# Differentially deregulated microRNAs contribute to ultraviolet radiation-induced photocarcinogenesis through immunomodulation: An-analysis of microRNAs expression profiling

**DOI:** 10.1101/2023.02.24.529976

**Authors:** Anshu Agarwal, Vikash Kansal, Humaira Farooqi, Vijay Kumar Singh, Ram Prasad

## Abstract

MicroRNAs (miRNAs) are short non-coding RNA molecules (18-25 nucleotides) that regulate several fundamental biological processes. Emerging evidence has shown more than 1500 miRNAs functions in the cell cycle, proliferation, apoptosis, oxidative stress, immune response, DNA damage, and epigenetics alterations. miRNAs are bidirectionally in nature and act as a tumor suppressor and as an oncogene through crosstalk between tumor cells and immune cells. Although the roles of miRNAs in several cancers are well studied, little is known about ultraviolet B (UVB) radiation-induced skin cancer. Here, we performed a comprehensive screening of 1281 miRNAs in tumor tissues and compared their expression with normal skin. Our results demonstrate that the expression levels of 587 miRNAs were altered in tumor tissues compared to their expression in normal skin. The expression of 337 miRNAs was upregulated from 1.5-12 folds, while the expression of 250 miRNAs was downregulated up to 1.5-10 folds in tumors. Further, intraperitoneal injection of a mimic of down-regulated miR-15b (30nM) and an inhibitor of upregulated miR-133a (20nM) protect UVB-induced suppression of contact hypersensitivity (CHS) response. In conclusion, we identified a network of altered miRNAs in tumors that can serve as prognostic biomarkers and therapeutic targets to manage photocarcinogenesis effectively.

## 1. Introduction

Skin, an important protective shield of the entire body, acts as a barrier to block the penetration of ultraviolet radiation (UV) and contact with foreign particles. Solar UV radiation is one of the critical risk factors that initiate carcinogenic events and promote skin cancer by inducing inflammatory responses, oxidative stress, immunosuppression, DNA damage, and gene mutations, all of which have been implicated in a variety of skin diseases, including the development of skin cancers [1–4]. UV radiation-induced inflammation plays a crucial role in all three stages of tumor development (initiation, promotion, and progression). Chronic exposure to UV radiation is a well-recognized etiological factor for all types of skin cancers, including basal cell carcinoma (BCC), squamous cell carcinoma (SCC), and melanoma, and which accounts for the approximately 1.3 million new cases of skin cancer each year in the USA [5]. The incidence of skin cancer is nearly equal to other malignancies in all other organs [6], thus representing a significant public health problem. Skin cancer development and progression is a complex process accompanied by multiple molecular changes at cellular levels. Despite intensive investigations, the precise mechanisms by which UVB radiation causes skin cancer remains unclear. Therefore, to uncover the molecular mechanisms involved in photocarcinogenesis, more novel makers must be discovered in this area.

MicroRNAs (miRNAs) are small non-coding RNA molecules (18-25 nucleotides) that play an essential role in various physiological functions in mammals and other multicellular organisms. A single miRNA targets up to hundreds of mRNAs. Approximately 30-60% of all human genes are affected by miRNA regulation. It has been reported that miRNAs affect several diseases and cancer-related processes such as proliferation, cell cycle control, apoptosis, differentiation, migration, and metabolism [7–10] and function as oncogenes or tumor-suppressor genes. This dual role of miRNAs has been reported in various studies. As tumor suppressors, they repress oncogenic targets but are usually downregulated in cancer tissues [11]. Others are upregulated and have a stimulating role in cancer progression [12, 13]. The biogenesis of miRNA is a three-step process; (i) transcription of primary miRNA (pri-miRNA) from the miRNA genes, (ii) partially processed precursor miRNA (pre-miRNA) in nuclei, and (iii) the generation of mature miRNAs into the cytoplasm (**Fig. 1**). Pri-miRNA is typically large transcripts and forms stem-loop structures.

**Figure 1:**
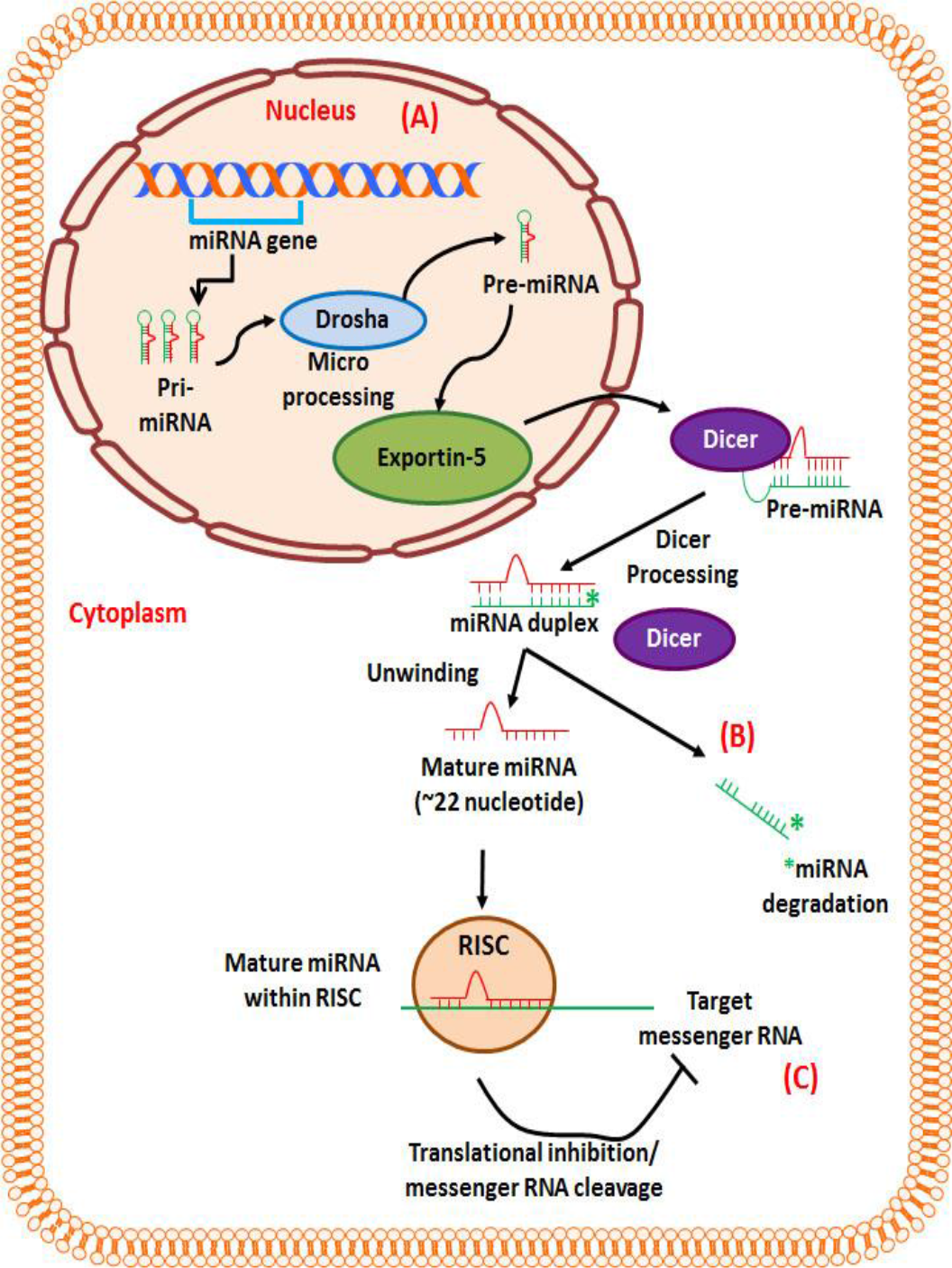
Biogenesis of miRNAs. **(A)** In the nucleus, ribonuclease enzyme II transcribes the pri-miRNA transcript. Drosha and its co factor Pasha form the microprocessor complex that cleaves pri-miRNA transcript into pre-miRNA which share a stem loop structure. These pre-miRNAs are transported by exportin-5 into cytoplasm from nucleus. **(B)** In the cytoplasm, Dicer unwind the miRNA duplex into mature miRNA and complementary strand (miRNA*, which rapidly degraded), and loaded mature miRNA into RNA-induced silencing complex (RISC). **(C)** Depending on the degree of complementarity between the target mRNA and the miRNA, the binding ether stops the translation by cleaving the target mRNA or suppress translation by binding to impeding ribosomal reading of the mRNA (incomplete complementarity). miRNA, microRNA; pri-miRNA, primary microRNA; pre-miRNA, precursor -microRNA; RISC, RNA- induced silencing complex.

Further, the stem-loop is cleaved off by the microprocessor machinery. Drosha (ribonuclease enzyme III) and its cofactor Pasha (DiGeorge syndrome critical region gene 8) form pre-miRNA (∼60-100 nucleotide). After successful cleavage, the pre-miRNA is bound by exportin-5 and exported from the nucleus to the cytoplasm [14, 15]. In the cytoplasm, pre-miRNA undergoes the next step of processing mediated by Dicer to produce the mature miRNA. The Dicer (ribonuclease enzyme III) cleaves RNAs into ∼22 nucleotide products [16, 17]. After cleavage, one strand of the miRNA duplex is preferentially incorporated into the RISC complex and binds to Argonaute (AGO) protein. The other strand, shown in **Fig. 1**, referred to as miRNA* (a complementary strand labeled by a star in the figure), normally degraded.

In some cases, miRNAs* can also be functional [18]. Although miRNAs are tiny in size but heavily involved in mammalian development through the regulation of various genes and represent a novel class of potential biomarkers or therapeutic targets. As a new layer of gene regulation mechanism, miRNAs have diverse functions, and deregulation alters normal cell growth and development, leading to various disorders, including human cancers. MiRNAs play a central role in immune regulation by modulating immune cells [19]. The expression of miRNAs has been demonstrated to be highly specific for tissues and developmental stages. Here to determine the effect of UVB radiation on miRNAs expression and their role in photocarcinogenesis, we performed a miRNAs expression profiling for more than 1000 miRNAs and provided a wide range of altered miRNAs expression in tumors.

## 2. Results

### 2.1. UVB radiation-induced alterations in miRNAs expression

First, to determine the effect of UVB radiation on miRNA expression in skin tumors, we performed miRNA array profiling using 7^th^-generation sequencing, which contains 3100 capture probes targeting human, mouse, and rat miRNAs registered in the miRBASE 18.0. These 3100 capture probes target 1281 mouse miRNAs (**Fig. 2A)**. Our miRNAs profiling revealed that the expression levels of 587 miRNAs were changed in UVB-induced tumors, while the levels of 694 miRNAs remained unchanged (**Fig. 2B)**. Out of 587 altered miRNAs, the expression levels of 250 miRNAs were downregulated, while the expression levels of 337 miRNAs were upregulated (**Fig. 2C)** in tumors.

**Figure 2:**
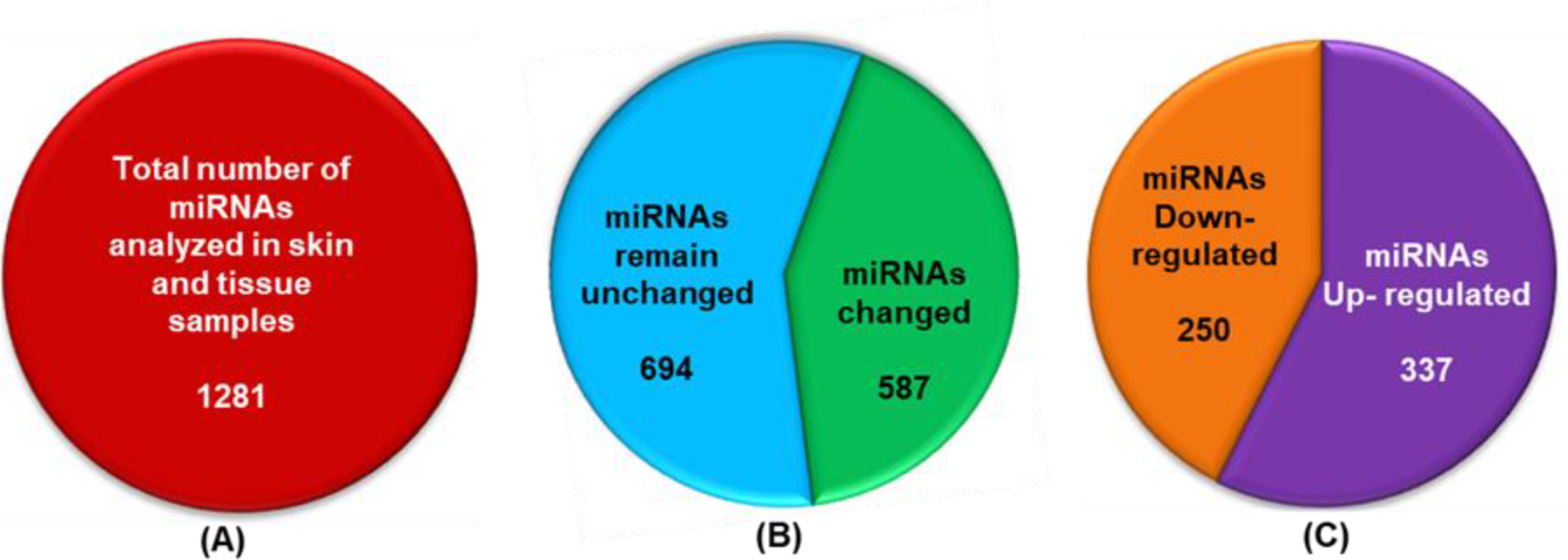
Total number of miRNAs profiled in tumors and normal skin of SKH-1 hairless mice. SKH-1 hairless mice were exposed to UVB radiation (180mJ/cm^2^) for up to 24 weeks, and tumor and skin tissue were harvested for miRNAs isolation using Trizol-chloroform extraction method. **(A)** Total number of miRNAs measured in tumors and skin tissues, **(B)** The number of altered miRNAs expression in tumors, **(C)**The number of downregulated, upregulated and aberrantly expressed miRNAs.

### 2.2. Down-regulation of miRNAs in UVB tumors

The loss of miRNA expression is closely related to tumor cell growth and has been implicated in tumorigenesis [20]. Therefore, we next identified the total number of down- regulated miRNAs in the tumor tissue. As shown in **Fig. 3A**, Out of 250 down-regulated miRNAs (**Table 1**), the expression of 165 miRNAs was reduced between 1.01-2.0 fold, while about 55 miRNAs’ expression was reduced between 2.01-4.0 fold in the tumors. The expression of 17 miRNAs was decreased within the range of 4.01-6.0 fold, and 13 miRNAs were downregulated more than 6 folds in the tumors compared to the normal skin. These down-regulated miRNAs may act as tumor suppressors.

**Figure 3:**
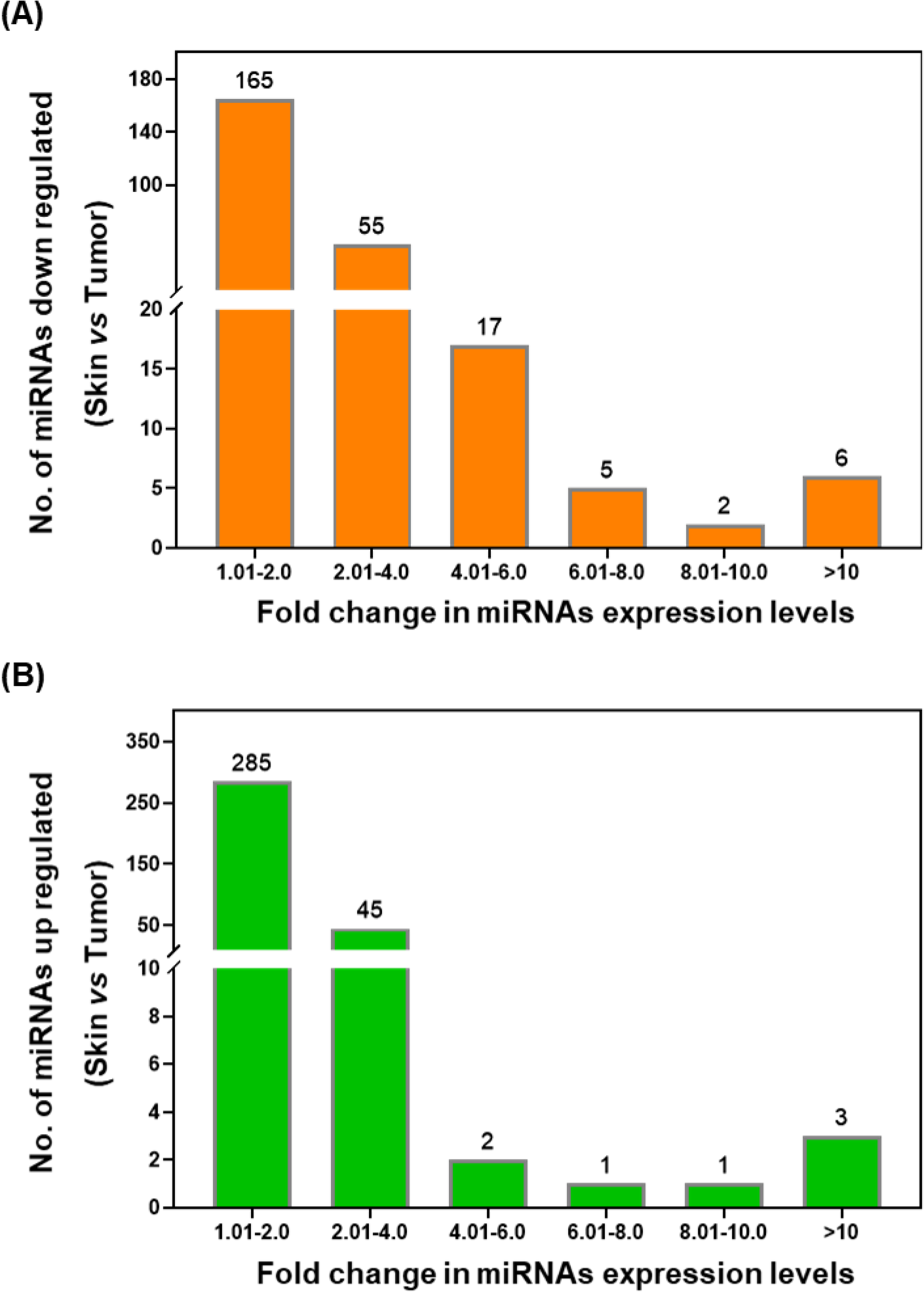
Graphical representation of fold change in the expression of miRNAs in UVB induced tumors. Graph showing difference in down regulated miRNAs **(A)** and upregulated miRNAs **(B).**

**Table 1:** Fold change expression of down-regulated miRNAs in UVB-induced tumors (UT). Data were compared with non-UVB exposed normal skin (CS).

| Probe ID | Annotation | AvgHy3 | UT | CS | logFC | Fold Change<br>(CS vs. UT) |
| --- | --- | --- | --- | --- | --- | --- |
| 42866 | mmu-miR-451a | 14.22 | 3.31 | -3.46 | -6.77 | -109.45 |
| 29802 | mmu-miR-144-3p | 11.65 | 3.57 | -2.79 | -6.36 | -82.18 |
| 32946 | mmu-miR-3107-5p/mmu-miR-486-5p | 9.44 | 3.02 | -1.21 | -4.23 | -18.75 |
| 169104 | mmu-miR-5099 | 12.49 | 2.80 | -1.12 | -3.91 | -15.05 |
| 169268 | mmu-miR-5112 | 8.49 | 2.45 | -1.07 | -3.52 | -11.48 |
| 17280 | mmu-miR-15b-5p | 11.39 | 2.70 | -0.71 | -3.41 | -10.66 |
| 148472 | mmu-miR-201-5p | 8.55 | 2.36 | -1.05 | -3.41 | -10.62 |
| 42682 | mmu-miR-25-3p | 9.25 | 2.64 | -0.66 | -3.30 | -9.84 |
| 46320 | mmu-miR-31-3p | 9.29 | 2.46 | -0.62 | -3.08 | -8.46 |
| 30687 | mmu-miR-93-5p | 10.13 | 2.50 | -0.47 | -2.97 | -7.85 |
| 10967 | mmu-miR-16-5p | 13.27 | 2.54 | -0.41 | -2.95 | -7.72 |
| 19599 | mmu-miR-106a-5p | 10.38 | 2.37 | -0.49 | -2.86 | -7.25 |
| 148578 | mmu-miR-541-3p | 10.32 | 2.38 | -0.44 | -2.82 | -7.07 |
| 169075 | mmu-miR-92a-3p | 9.63 | 2.18 | -0.50 | -2.68 | -6.40 |
| 11052 | mmu-miR-31-5p | 10.26 | 2.23 | -0.31 | -2.54 | -5.81 |
| **145845** | mmu-miR-20a-5p | 10.97 | 2.06 | -0.47 | -2.54 | -5.80 |
| **148344** | mmu-miR-669l-3p | 12.23 | 2.10 | -0.39 | -2.49 | -5.61 |
| **42640** | mmu-miR-20b-5p | 9.30 | 1.96 | -0.49 | -2.45 | -5.45 |
| **169336** | mmu-miR-17-5p | 10.42 | 2.04 | -0.40 | -2.44 | -5.43 |
| **10947** | mmu-miR-142-3p | 12.53 | 1.96 | -0.39 | -2.36 | -5.12 |
| **10997** | mmu-miR-19a-3p | 9.96 | 1.88 | -0.46 | -2.34 | -5.06 |
| **42630** | mmu-miR-140-3p | 9.88 | 2.09 | -0.24 | -2.32 | -5.01 |
| **148101** | mmu-miR-669d-2-3p/mmu-miR-669d-3p | 12.42 | 1.98 | -0.33 | -2.31 | -4.96 |
| **19603** | SNORD13 | 9.85 | 2.00 | -0.31 | -2.31 | -4.96 |
| **27720** | mmu-miR-15a-5p | 12.24 | 2.15 | -0.14 | -2.29 | -4.91 |
| **19582** | mmu-miR-106b-5p | 11.34 | 1.99 | -0.26 | -2.25 | -4.75 |
| **146164** | mmu-miR-1958 | 11.82 | 1.95 | -0.16 | -2.12 | -4.34 |
| **146155** | mmu-miR-2137 | 12.40 | 1.96 | -0.15 | -2.12 | -4.33 |
| **14302** | mmu-miR-374b-5p/mmu-miR-374c-5p | 7.96 | 1.83 | -0.28 | -2.11 | -4.32 |
| **147506** | mmu-miR-21a-5p | 13.80 | 1.25 | -0.80 | -2.04 | -4.12 |
| **46204** | SNORD38B | 9.68 | 1.88 | -0.12 | -2.01 | -4.01 |
| **145852** | mmu-miR-210-3p | 9.08 | 1.89 | -0.07 | -1.96 | -3.89 |
| **18739** | mmu-miR-186-5p | 7.64 | 1.92 | -0.03 | -1.95 | -3.87 |
| **13143** | mmu-miR-301a-3p | 8.53 | 1.87 | -0.04 | -1.91 | -3.76 |
| **14290** | mmu-miR-541-5p | 8.20 | 1.53 | -0.37 | -1.90 | -3.72 |
| **13147** | mmu-miR-96-5p | 8.59 | 1.80 | -0.09 | -1.89 | -3.70 |
| **168556** | mmu-miR-3096b-5p | 9.50 | 1.50 | -0.35 | -1.86 | -3.62 |
| **11024** | mmu-miR-223-3p | 11.55 | 1.80 | -0.04 | -1.85 | -3.59 |
| **168706** | mmu-miR-5129-5p | 8.60 | 1.33 | -0.50 | -1.84 | -3.57 |
| **148233** | mmu-miR-3096a-3p | 10.79 | 1.38 | -0.44 | -1.82 | -3.53 |
| **11245** | mmu-miR-433-5p | 10.60 | 1.48 | -0.33 | -1.81 | -3.51 |
| **168880** | mmu-miR-489-5p | 7.39 | 1.16 | -0.65 | -1.81 | -3.51 |
| **14289** | mmu-miR-540-3p | 9.68 | 1.43 | -0.38 | -1.81 | -3.50 |
| **148098** | mmu-miR-374b-5p | 8.54 | 1.70 | -0.09 | -1.79 | -3.46 |
| **148238** | mmu-miR-3096a-5p | 9.47 | 1.59 | -0.18 | -1.78 | -3.43 |
| **148528** | mmu-miR-196a-1-3p | 8.40 | 1.15 | -0.63 | -1.78 | -3.43 |
| **145798** | mmu-miR-142-5p | 10.46 | 1.55 | -0.22 | -1.78 | -3.43 |
| **148087** | mmu-miR-669d-2-3p | 9.70 | 1.63 | -0.03 | -1.66 | -3.16 |
| **146192** | mmu-miR-669m-3p | 11.14 | 1.75 | 0.10 | -1.65 | -3.14 |
| **10998** | mmu-miR-19b-3p | 10.88 | 1.55 | -0.10 | -1.65 | -3.13 |
| **42811** | mmu-miR-542-5p | 8.23 | 1.68 | 0.07 | -1.61 | -3.04 |
| **148035** | mmu-miR-3084-5p | 9.95 | 1.55 | -0.04 | -1.59 | -3.00 |
| **169340** | mmu-miR-3096a-3p/mm-miR-3096b-3p | 9.99 | 1.00 | -0.53 | -1.53 | -2.88 |
| **146012** | mmu-miR-1949 | 9.75 | 0.93 | -0.52 | -1.44 | -2.72 |
| **145633** | mmu-let-7d-3p | 8.99 | 1.36 | -0.07 | -1.43 | -2.69 |
| **46917** | mmu-miR-205-5p | 13.85 | 1.41 | 0.01 | -1.40 | -2.64 |
| **4700** | mmu-miR-140-5p | 8.84 | 1.41 | 0.02 | -1.40 | -2.64 |
| **17608** | mmu-miR-425-5p | 7.84 | 1.30 | -0.10 | -1.40 | -2.63 |
| **17427** | mmu-miR-200c-3p | 10.57 | 1.42 | 0.02 | -1.40 | -2.63 |
| **19585** | mmu-miR-148b-3p | 8.26 | 1.43 | 0.06 | -1.37 | -2.59 |
| **27558** | mmu-miR-155-5p | 7.66 | 1.31 | -0.02 | -1.33 | -2.52 |
| **169347** | mmu-miR-5622-5p | 9.16 | 1.22 | -0.05 | -1.26 | -2.40 |
| **42702** | mmu-miR-30c-1-3p | 7.63 | 1.24 | -0.02 | -1.26 | -2.40 |
| **17610** | mmu-miR-677-5p | 8.36 | 1.27 | 0.02 | -1.25 | -2.38 |
| **10975** | mmu-miR-182-5p | 8.49 | 1.17 | -0.07 | -1.24 | -2.36 |
| **42919** | mmu-miR-203-5p | 8.96 | 1.40 | 0.16 | -1.23 | -2.35 |
| **148484** | mmu-miR-3084-3p | 11.53 | 1.02 | -0.20 | -1.22 | -2.33 |
| **9938** | mmu-let-7i-5p | 11.74 | 1.21 | -0.01 | -1.21 | -2.32 |
| **10946** | mmu-miR-141-3p | 10.95 | 1.17 | -0.03 | -1.20 | -2.30 |
| **19007** | SNORD3@ | 12.94 | 0.93 | -0.26 | -1.20 | -2.30 |
| **148676** | mmu-miR-1186b | 10.48 | 0.92 | -0.28 | -1.20 | -2.29 |
| **169277** | mmu-miR-3970 | 9.60 | 1.06 | -0.12 | -1.17 | -2.26 |
| **145661** | SNORD65 | 8.08 | 1.06 | -0.09 | -1.15 | -2.22 |
| **27533** | mmu-miR-320-3p | 8.77 | 1.31 | 0.16 | -1.15 | -2.22 |
| **46438** | mmu-let-7g-5p | 11.41 | 1.35 | 0.21 | -1.14 | -2.20 |
| **42730** | mmu-miR-423-3p | 8.51 | 0.94 | -0.18 | -1.12 | -2.17 |
| **29490** | mmu-miR-7a-5p | 10.34 | 1.00 | -0.10 | -1.09 | -2.13 |
| **42571** | mmu-miR-129-1-3p | 7.53 | 0.91 | -0.17 | -1.08 | -2.11 |
| **168966** | mmu-miR-28a-5p/mmu-miR-28c | 8.67 | 1.01 | -0.06 | -1.07 | -2.10 |
| **168872** | mmu-miR-24-1-5p | 9.44 | 1.14 | 0.07 | -1.06 | -2.09 |
| **11022** | mmu-miR-221-3p | 9.36 | 1.08 | 0.03 | -1.06 | -2.08 |
| **31388** | mmu-miR-291a-5p | 10.55 | 0.73 | -0.31 | -1.05 | -2.07 |
| **17854** | mmu-miR-106b-3p | 8.53 | 1.18 | 0.14 | -1.04 | -2.06 |
| **168859** | mmu-miR-3962 | 10.72 | 1.20 | 0.17 | -1.03 | -2.04 |
| **10985** | mmu-miR-191-5p | 9.98 | 1.25 | 0.22 | -1.03 | -2.04 |
| **29190** | mmu-miR-708-5p | 8.99 | 1.28 | 0.27 | -1.00 | -2.01 |
| **42707** | mmu-miR-294-5p | 12.23 | 0.57 | -0.43 | -1.00 | -2.00 |
| **146004** | mmu-miR-2136 | 7.47 | 1.12 | 0.13 | -0.99 | -1.99 |
| **148051** | mmu-miR-770-3p | 7.89 | 0.97 | -0.01 | -0.97 | -1.96 |
| **146111** | mmu-miR-767 | 8.42 | 0.85 | -0.12 | -0.97 | -1.96 |
| **147199** | mmu-miR-27b-3p | 11.90 | 0.91 | -0.05 | -0.96 | -1.94 |
| **28547** | mmu-miR-675-5p | 9.89 | 0.94 | 0.00 | -0.95 | -1.93 |
| **11184** | mmu-miR-99b-5p | 9.63 | 0.95 | 0.00 | -0.94 | -1.92 |
| **11023** | mmu-miR-222-3p | 9.70 | 0.92 | -0.02 | -0.94 | -1.92 |
| **148416** | mmu-miR-3102-5p | 7.87 | 0.68 | -0.26 | -0.93 | -1.91 |
| **169330** | mmu-miR-23b-3p | 13.09 | 1.01 | 0.12 | -0.89 | -1.86 |
| **28450** | mmu-miR-291b-5p | 9.91 | 0.72 | -0.17 | -0.89 | -1.85 |
| **145663** | SNORD68 | 11.57 | 0.92 | 0.03 | -0.89 | -1.85 |
| **11260** | mmu-miR-151-5p | 8.24 | 1.04 | 0.16 | -0.88 | -1.84 |
| **168630** | mmu-miR-5121 | 9.19 | 0.57 | -0.30 | -0.87 | -1.83 |
| **17752** | mmu-let-7f-5p | 9.95 | 1.18 | 0.31 | -0.87 | -1.83 |
| **10990** | mmu-miR-196a-5p | 7.51 | 0.40 | -0.45 | -0.86 | -1.81 |
| **145666** | SNORD110 | 8.73 | 0.83 | -0.02 | -0.85 | -1.81 |
| **19008** | SNORD2 | 12.27 | 0.61 | -0.24 | -0.84 | -1.79 |
| **146199** | mmu-miR-1961 | 8.41 | 1.05 | 0.21 | -0.84 | -1.79 |
| **146112** | mmu-miR-30b-5p | 11.23 | 1.15 | 0.31 | -0.83 | -1.78 |
| **11256** | mmu-miR-470-5p | 9.04 | 0.79 | -0.04 | -0.83 | -1.78 |
| **145968** | mmu-let-7d-5p | 11.44 | 1.02 | 0.19 | -0.83 | -1.77 |
| **27575** | mmu-miR-711 | 9.56 | 1.17 | 0.35 | -0.82 | -1.77 |
| **148523** | mghv-miR-M1-8-3p | 11.82 | 0.50 | -0.31 | -0.81 | -1.75 |
| **10988** | mmu-miR-194-5p | 7.42 | 0.72 | -0.08 | -0.80 | -1.74 |
| **10936** | mmu-miR-130b-3p | 7.66 | 0.69 | -0.11 | -0.80 | -1.74 |
| **146087** | mmu-miR-1894-3p | 8.36 | 0.88 | 0.08 | -0.80 | -1.74 |
| **30831** | mmu-miR-804 | 7.40 | 1.21 | 0.42 | -0.79 | -1.73 |
| **17506** | mmu-miR-24-3p | 13.49 | 1.04 | 0.25 | -0.78 | -1.72 |
| **147162** | mmu-let-7a-5p | 11.08 | 0.98 | 0.20 | -0.78 | -1.72 |
| **17478** | mmu-miR-429-3p | 8.89 | 1.14 | 0.37 | -0.78 | -1.71 |
| **146172** | mmu-miR-1892 | 8.93 | 0.65 | -0.12 | -0.77 | -1.70 |
| **42953** | mmu-miR-101b-3p | 9.31 | 0.99 | 0.23 | -0.76 | -1.69 |
| **17597** | mmu-miR-467b-3p | 9.91 | 1.00 | 0.25 | -0.74 | -1.67 |
| **148100** | mmu-miR-1947-3p | 12.96 | 0.37 | -0.36 | -0.73 | -1.66 |
| **145859** | mmu-miR-33-5p | 9.39 | 0.79 | 0.06 | -0.73 | -1.65 |
| **11020** | mmu-miR-22-3p | 12.57 | 0.93 | 0.21 | -0.72 | -1.65 |
| **28966** | mmu-miR-574-3p | 8.91 | 0.73 | 0.02 | -0.71 | -1.64 |
| **42923** | mmu-miR-30c-5p | 11.58 | 0.95 | 0.26 | -0.69 | -1.61 |
| **148107** | mmu-miR-3104-3p | 7.93 | 0.92 | 0.23 | -0.69 | -1.61 |
| **168824** | mmu-miR-5100 | 15.78 | 0.89 | 0.20 | -0.68 | -1.61 |
| **168842** | mmu-miR-5105 | 7.38 | 0.68 | 0.01 | -0.67 | -1.59 |
| **28191** | mmu-miR-30e-5p | 10.73 | 0.95 | 0.28 | -0.67 | -1.59 |
| **145838** | mmu-miR-125b-1-3p | 8.42 | 0.74 | 0.07 | -0.67 | -1.59 |
| **148362** | mmu-miR-592-3p | 9.08 | 1.06 | 0.40 | -0.66 | -1.58 |
| **42605** | mmu-miR-503-3p | 11.29 | 0.54 | -0.12 | -0.66 | -1.58 |
| **168819** | mmu-miR-200a-3p | 10.35 | 0.87 | 0.22 | -0.65 | -1.57 |
| **145753** | mmu-miR-484 | 7.61 | 0.82 | 0.18 | -0.64 | -1.56 |
| **42827** | mmu-miR-652-3p | 8.04 | 0.75 | 0.13 | -0.62 | -1.54 |
| **145897** | mmu-miR-92b-3p | 8.50 | 0.23 | -0.39 | -0.62 | -1.53 |
| **169190** | mmu-miR-5117-3p | 11.20 | 0.73 | 0.11 | -0.61 | -1.53 |
| **169105** | mmu-miR-3963 | 15.95 | 1.14 | 0.53 | -0.61 | -1.53 |
| **11053** | mmu-miR-32-5p | 8.82 | 0.78 | 0.18 | -0.60 | -1.52 |
| **145993** | mmu-miR-1899 | 8.41 | 0.60 | 0.00 | -0.59 | -1.51 |
| **19600** | mmu-miR-17-3p | 9.34 | 0.41 | -0.17 | -0.59 | -1.50 |
| **11215** | mmu-miR-292-3p | 7.57 | 1.95 | 1.37 | -0.59 | -1.50 |
| **145701** | mmu-miR-668-3p | 7.90 | 0.81 | 0.23 | -0.58 | -1.49 |
| **14301** | mmu-miR-361-5p | 9.27 | 0.80 | 0.22 | -0.58 | -1.49 |
| **14285** | mmu-miR-487b-3p | 8.04 | 0.56 | -0.01 | -0.58 | -1.49 |
| **148059** | mmu-miR-493-5p | 8.92 | 0.24 | -0.33 | -0.57 | -1.48 |
| **169248** | mmu-miR-5108 | 7.39 | 0.40 | -0.16 | -0.57 | -1.48 |
| **169250** | mmu-miR-5109 | 13.60 | 0.69 | 0.13 | -0.56 | -1.48 |
| **148197** | mmu-miR-3081-5p | 8.34 | 0.46 | -0.10 | -0.56 | -1.47 |
| **42888** | mmu-miR-875-3p | 12.08 | 0.47 | -0.09 | -0.56 | -1.47 |
| **168708** | mmu-miR-296-5p | 7.47 | 0.78 | 0.22 | -0.56 | -1.47 |
| **11227** | mmu-miR-329-3p | 9.38 | 0.98 | 0.43 | -0.54 | -1.46 |
| **46210** | mmu-miR-1249-3p | 7.70 | 1.14 | 0.61 | -0.53 | -1.44 |
| **169246** | mmu-miR-5618-5p | 7.82 | 0.47 | -0.05 | -0.52 | -1.44 |
| **17489** | mmu-miR-710 | 10.34 | 0.54 | 0.03 | -0.51 | -1.43 |
| **30768** | mmu-miR-674-5p | 9.18 | 0.57 | 0.07 | -0.50 | -1.42 |
| **10138** | mmu-miR-130a-3p | 10.45 | 0.80 | 0.31 | -0.49 | -1.40 |
| **19606** | SNORD12 | 8.89 | 0.30 | -0.18 | -0.48 | -1.39 |
| **168807** | mmu-miR-3473c | 7.55 | 0.59 | 0.12 | -0.47 | -1.39 |
| **10955** | mmu-miR-148a-3p | 8.03 | 0.50 | 0.04 | -0.46 | -1.38 |
| **169374** | mmu-miR-184-5p | 11.08 | 0.34 | -0.12 | -0.46 | -1.38 |
| **148261** | mmu-miR-208a-5p | 7.23 | 0.46 | 0.00 | -0.46 | -1.38 |
| **168762** | mmu-miR-3964 | 7.99 | 0.18 | -0.27 | -0.45 | -1.37 |
| **11182** | mmu-miR-98-5p | 10.38 | 0.77 | 0.34 | -0.44 | -1.35 |
| **11234** | mmu-miR-350-3p | 7.72 | 0.57 | 0.14 | -0.44 | -1.35 |
| **168586** | mmu-miR-34a-5p | 9.78 | 0.11 | -0.32 | -0.43 | -1.35 |
| **46483** | mmu-miR-27a-3p | 11.82 | 0.44 | 0.01 | -0.43 | -1.34 |
| **147186** | mmu-miR-200b-3p | 10.53 | 0.67 | 0.27 | -0.41 | -1.33 |
| **46636** | mcmv-miR-M23-1-5p | 8.00 | -0.09 | -0.49 | -0.39 | -1.31 |
| **13140** | mmu-miR-138-5p | 8.44 | 0.41 | 0.04 | -0.37 | -1.29 |
| **148654** | mmu-miR-184-3p | 7.29 | 0.47 | 0.10 | -0.37 | -1.29 |
| **10986** | mmu-miR-193a-3p | 9.90 | 0.60 | 0.23 | -0.37 | -1.29 |
| **21498** | mmu-miR-654-3p | 7.21 | 0.63 | 0.26 | -0.36 | -1.29 |
| **19605** | SNORD6 | 7.44 | 0.69 | 0.34 | -0.34 | -1.27 |
| **148046** | mmu-miR-344-5p | 7.42 | -0.03 | -0.37 | -0.34 | -1.27 |
| **42744** | mmu-miR-23a-3p | 13.14 | 0.65 | 0.32 | -0.33 | -1.26 |
| **42636** | mmu-miR-28a-3p | 7.39 | 0.48 | 0.16 | -0.32 | -1.25 |
| **148391** | mmu-miR-3068-5p | 8.04 | 0.07 | -0.25 | -0.31 | -1.24 |
| **148424** | mmu-miR-201-3p | 8.42 | 0.14 | -0.17 | -0.31 | -1.24 |
| **46251** | mmu-miR-1193-3p | 7.35 | 0.03 | -0.28 | -0.31 | -1.24 |
| **42724** | mmu-miR-34b-3p | 9.92 | 0.23 | -0.08 | -0.30 | -1.24 |
| **145643** | mmu-miR-382-5p | 7.96 | 0.47 | 0.16 | -0.30 | -1.24 |
| **17810** | mmu-miR-29b-1-5p | 9.05 | 0.38 | 0.07 | -0.30 | -1.23 |
| **146065** | mmu-miR-1927 | 7.27 | 0.35 | 0.05 | -0.30 | -1.23 |
| **148218** | mghv-miR-M1-11-3p | 7.84 | 0.42 | 0.13 | -0.29 | -1.22 |
| **148485** | mghv-miR-M1-12-3p | 8.22 | 0.40 | 0.11 | -0.29 | -1.22 |
| **146106** | mmu-miR-1931 | 8.20 | 0.19 | -0.10 | -0.29 | -1.22 |
| **42494** | mmu-miR-712-3p | 7.58 | 0.27 | -0.02 | -0.29 | -1.22 |
| **42452** | mmu-miR-141-5p | 7.69 | 0.33 | 0.05 | -0.28 | -1.22 |
| **148166** | mmu-miR-3069-3p | 8.84 | 0.28 | 0.00 | -0.28 | -1.21 |
| **42879** | mmu-miR-92a-2-5p | 8.29 | 0.65 | 0.38 | -0.27 | -1.21 |
| **42488** | mmu-miR-466h-5p | 7.32 | 0.92 | 0.66 | -0.26 | -1.20 |
| **14328** | mmu-miR-124-3p | 7.45 | 0.50 | 0.24 | -0.25 | -1.19 |
| **148470** | mmu-miR-1264-3p | 10.00 | 0.12 | -0.13 | -0.25 | -1.19 |
| **148370** | mmu-miR-466n-3p | 9.63 | 0.22 | -0.02 | -0.24 | -1.18 |
| **10919** | mmu-miR-103-3p | 10.69 | 0.57 | 0.32 | -0.24 | -1.18 |
| **148172** | mmu-miR-216a-3p | 8.11 | -0.10 | -0.32 | -0.23 | -1.17 |
| **42570** | mmu-miR-194-2-3p | 7.35 | 0.33 | 0.11 | -0.22 | -1.16 |
| **42769** | mmu-let-7b-3p | 7.38 | 0.21 | 0.01 | -0.20 | -1.15 |
| **29852** | mmu-miR-9-3p | 7.32 | 0.08 | -0.12 | -0.19 | -1.14 |
| **145676** | mmu-miR-30e-3p | 8.54 | 0.50 | 0.31 | -0.19 | -1.14 |
| **13150** | mmu-miR-322-5p | 8.43 | 0.36 | 0.17 | -0.19 | -1.14 |
| **146133** | mmu-miR-1936 | 8.27 | 0.10 | -0.09 | -0.18 | -1.14 |
| **19013** | SNORD14B | 7.68 | 0.42 | 0.24 | -0.18 | -1.13 |
| **168566** | mmu-miR-5625-3p | 8.24 | -0.29 | -0.46 | -0.17 | -1.13 |
| **145638** | mmu-miR-29a-5p | 8.21 | 0.47 | 0.30 | -0.17 | -1.12 |
| **148179** | mmu-miR-3095-3p | 10.46 | 0.04 | -0.12 | -0.16 | -1.12 |
| **168687** | mmu-miR-29a-3p | 12.14 | 0.29 | 0.12 | -0.16 | -1.12 |
| **168662** | mmu-miR-5132-5p | 8.87 | -0.01 | -0.17 | -0.16 | -1.12 |
| **146086** | mmu-miR-30a-5p | 10.06 | 0.62 | 0.47 | -0.15 | -1.11 |
| **27536** | mmu-miR-190a-5p | 7.52 | 0.87 | 0.73 | -0.15 | -1.11 |
| **17953** | mmu-miR-183-3p | 10.40 | 0.07 | -0.08 | -0.15 | -1.11 |
| **145640** | mmu-miR-328-3p | 8.53 | 0.02 | -0.13 | -0.14 | -1.10 |
| **33114** | mmu-miR-455-3p | 7.46 | 0.05 | -0.09 | -0.14 | -1.10 |
| **17273** | mg hv-miR-M1-6-3p | 8.28 | 0.67 | 0.53 | -0.14 | -1.10 |
| **28019** | mmu-miR-10a-3p | 7.92 | 0.14 | 0.01 | -0.13 | -1.09 |
| **148242** | mmu-miR-205-3p | 8.63 | -0.14 | -0.27 | -0.13 | -1.09 |
| **148121** | mmu-miR-155-3p | 9.32 | 0.18 | 0.05 | -0.13 | -1.09 |
| **11251** | mmu-miR-465a-5p | 7.75 | 0.68 | 0.56 | -0.13 | -1.09 |
| **168968** | mmu-miR-147-3p | 8.94 | 0.27 | 0.15 | -0.12 | -1.09 |
| **147283** | mmu-miR-137-5p | 8.05 | 0.50 | 0.39 | -0.12 | -1.08 |
| **19596** | mmu-miR-30d-5p | 9.79 | 0.59 | 0.47 | -0.12 | -1.08 |
| **17818** | mmu-miR-27a-5p | 7.25 | 0.30 | 0.18 | -0.11 | -1.08 |
| **148546** | mmu-miR-500-5p | 7.30 | 0.33 | 0.21 | -0.11 | -1.08 |
| **146008** | mmu-miR-26b-5p | 11.47 | 0.55 | 0.43 | -0.11 | -1.08 |
| **148646** | mmu-miR-467a-3p | 11.52 | 0.27 | 0.16 | -0.11 | -1.08 |
| **168797** | mmu-miR-3968 | 11.65 | -0.28 | -0.38 | -0.10 | -1.07 |
| **42808** | mmu-miR-874-3p | 7.44 | -0.12 | -0.22 | -0.10 | -1.07 |
| **42738** | mmu-miR-340-3p | 7.65 | 0.19 | 0.10 | -0.09 | -1.07 |
| **10923** | mmu-miR-107-3p | 10.15 | 0.39 | 0.30 | -0.09 | -1.06 |
| **42950** | mmu-miR-24-2-5p | 9.57 | 0.38 | 0.30 | -0.09 | -1.06 |
| **11040** | mmu-miR-29b-3p | 11.90 | 0.32 | 0.24 | -0.08 | -1.06 |
| **17825** | mmu-miR-338-5p | 7.72 | 0.13 | 0.05 | -0.08 | -1.06 |
| **42532** | mmu-miR-22-5p | 9.41 | 0.45 | 0.38 | -0.08 | -1.05 |
| **148225** | mmu-miR-3102-5p.2-5p | 7.38 | -0.03 | -0.09 | -0.07 | -1.05 |
| **10977** | mmu-miR-183-5p | 9.22 | -0.11 | -0.17 | -0.06 | -1.04 |
| **14272** | mmu-miR-542-3p | 7.86 | 0.34 | 0.28 | -0.06 | -1.04 |
| **42464** | mghv-miR-M1-2-3p | 9.14 | 0.67 | 0.62 | -0.05 | -1.04 |
| **148128** | mmu-miR-3090-5p | 9.16 | -0.10 | -0.15 | -0.05 | -1.03 |
| **11078** | mmu-miR-365-3p | 9.43 | 0.42 | 0.37 | -0.04 | -1.03 |
| **16681** | mmu-miR-721 | 9.22 | -0.05 | -0.09 | -0.04 | -1.02 |
| **30787** | mmu-miR-125b-5p | 12.75 | 0.37 | 0.34 | -0.03 | -1.02 |
| **148689** | mmu-miR-3099-5p | 9.18 | 0.18 | 0.15 | -0.03 | -1.02 |
| **42929** | mmu-miR-25-5p | 10.96 | 0.71 | 0.69 | -0.03 | -1.02 |
| **17942** | mmu-miR-125a-3p | 7.67 | 0.02 | -0.01 | -0.03 | -1.02 |
| **42804** | mmu-miR-712-5p | 8.16 | 0.00 | -0.02 | -0.02 | -1.02 |
| **46292** | mmu-miR-5097 | 11.74 | -0.68 | -0.69 | -0.02 | -1.01 |
| **148303** | mmu-miR-3106-5p | 7.42 | -0.26 | -0.27 | -0.02 | -1.01 |
| **169060** | mmu-miR-3961 | 12.40 | -0.25 | -0.27 | -0.01 | -1.01 |
| **148270** | mmu-miR-669b-3p | 11.78 | 0.28 | 0.27 | -0.01 | -1.01 |
| **145678** | mmu-miR-150-5p | 9.57 | 0.11 | 0.10 | -0.01 | -1.01 |

### 2.3. Upregulation of miRNAs in UVB tumors

UV radiation causes several mutations in gene status to induce tumors, such as the p53 mutation regulated by miRNAs. In our miRNAs profiling results, we observed that the levels of 3377 miRNAs were upregulated in tumors (**Fig. 3B**, **Table 2**). In the tumor tissue, out of 337 upregulated miRNAs, the expression of 285 miRNAs was increased up to 2.0 fold, while about 45 miRNAs’ expression was increased between 2.01-4.0 fold. The expression of 7 miRNAs was highly upregulated more than 6 folds in the UVB-induced tumors compared with their expression in the normal skin. These upregulated miRNAs may act as an oncogene to promote photocarcinogenesis.

**Table 2:**
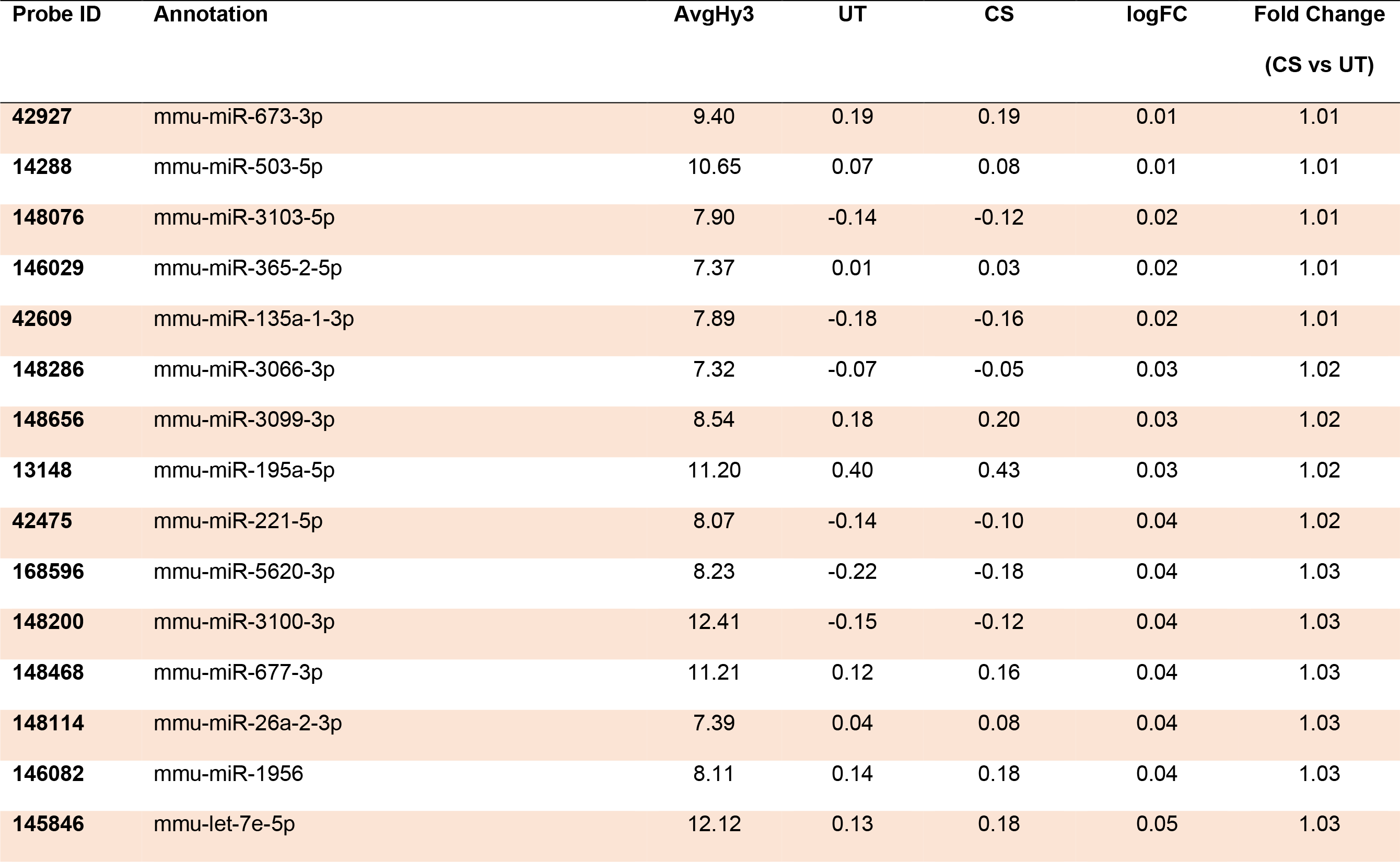
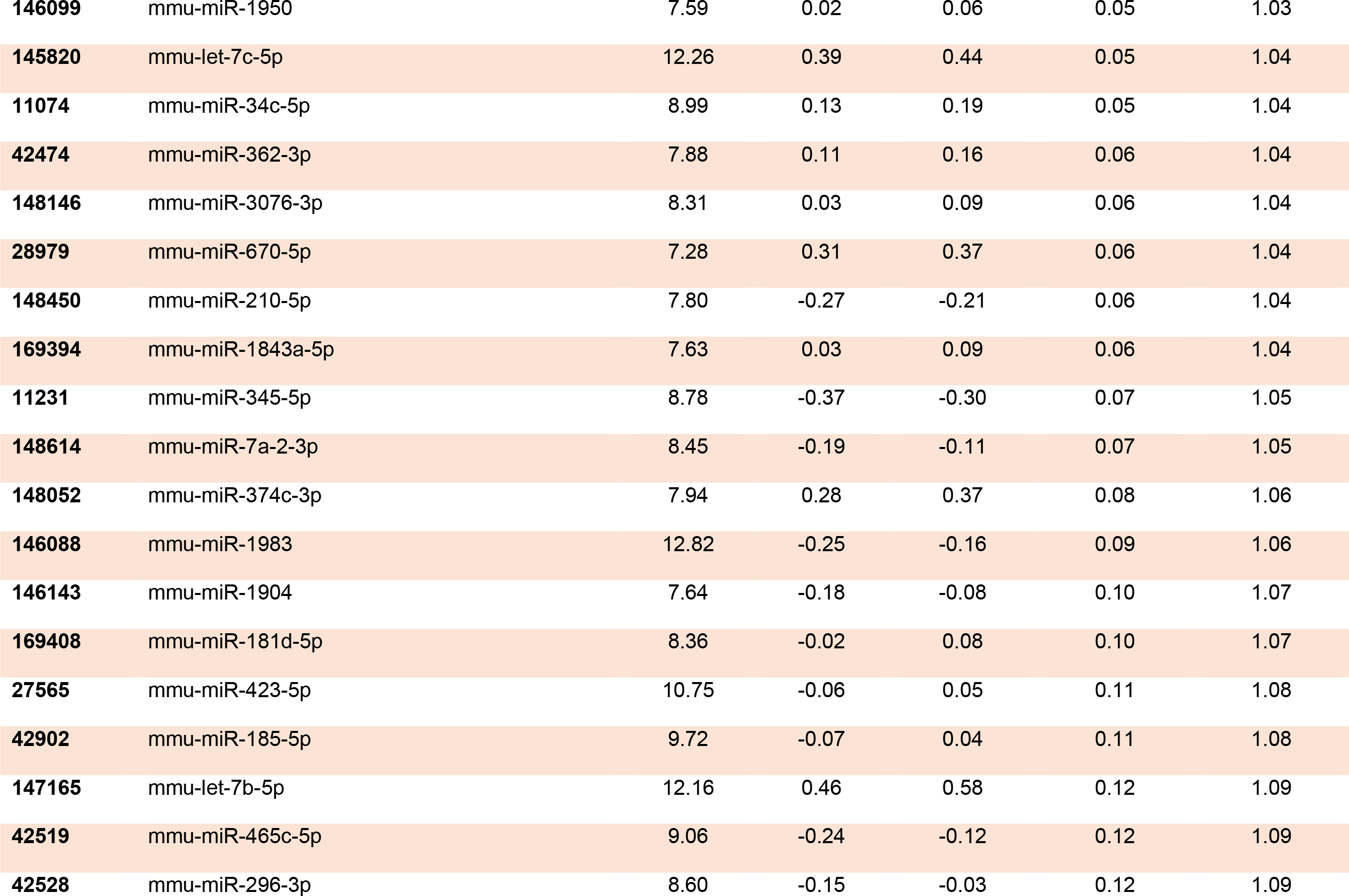
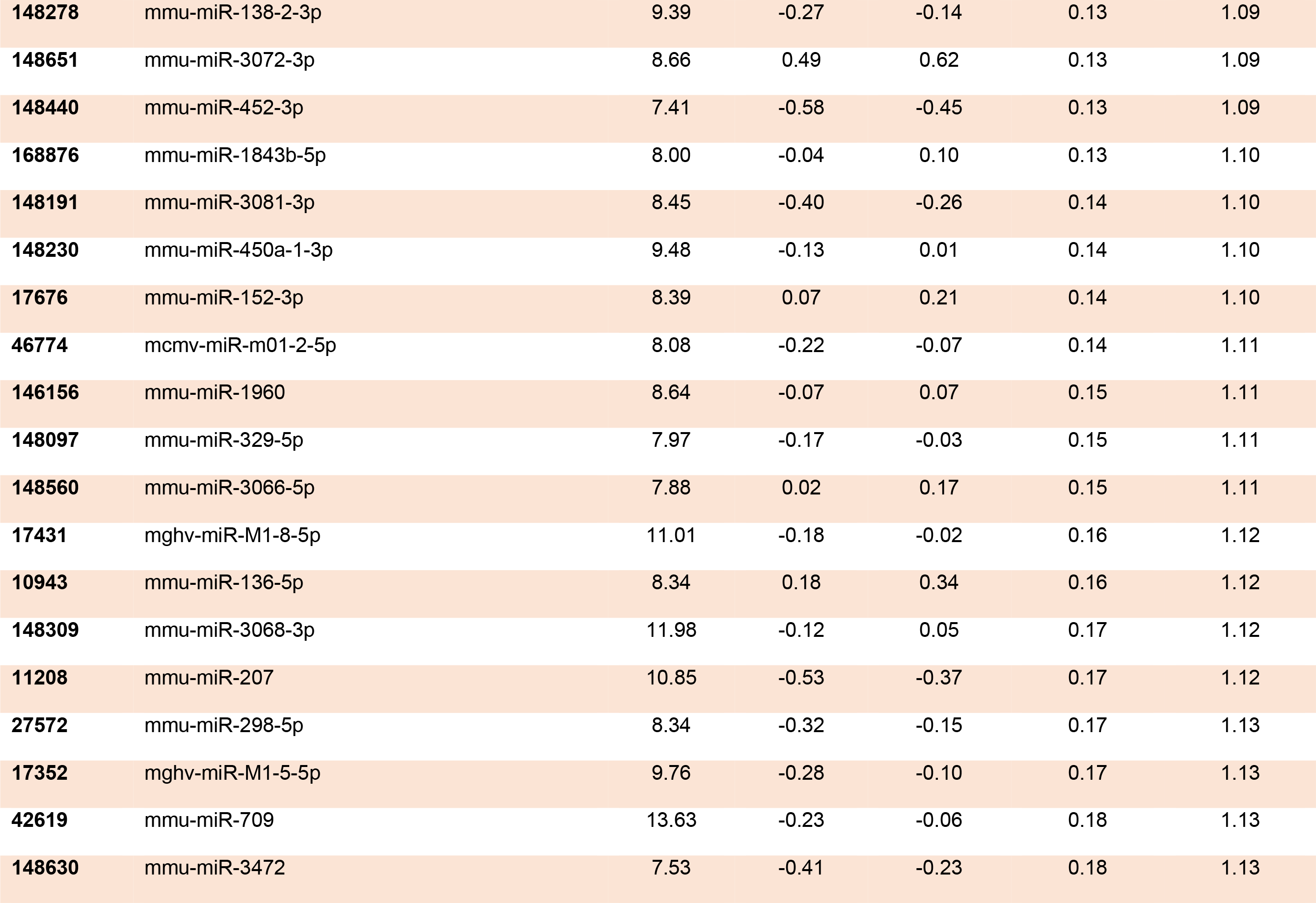
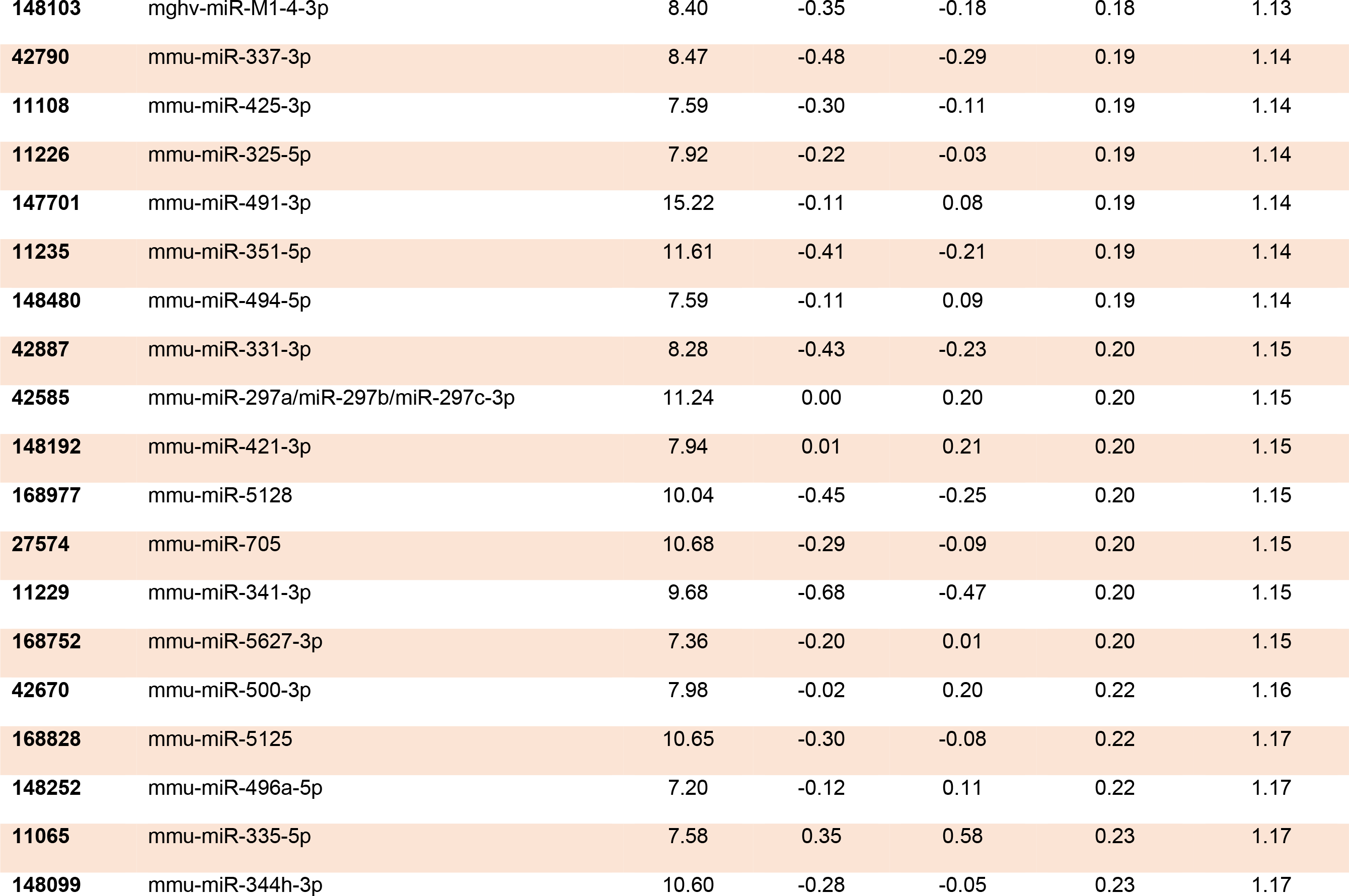
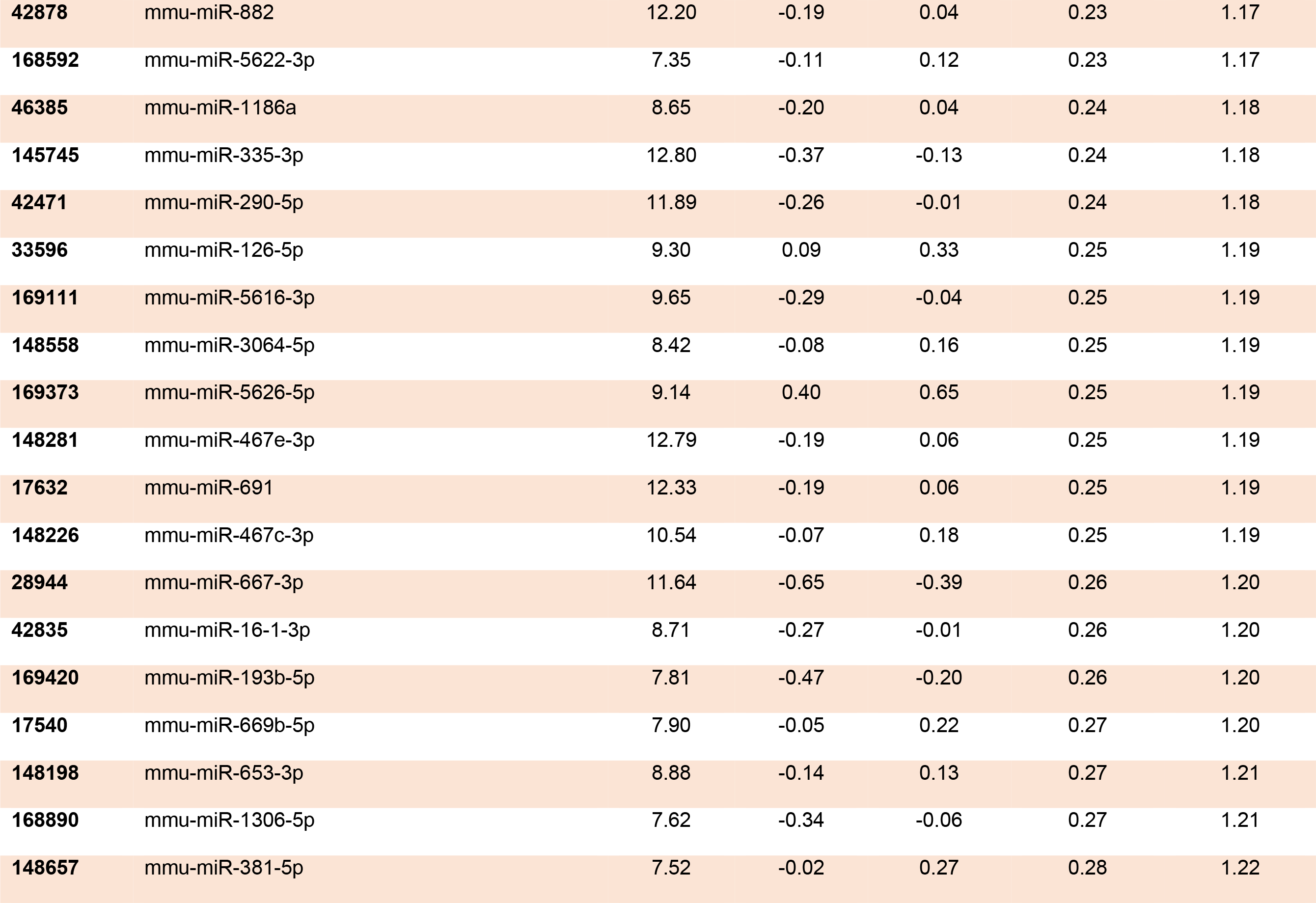
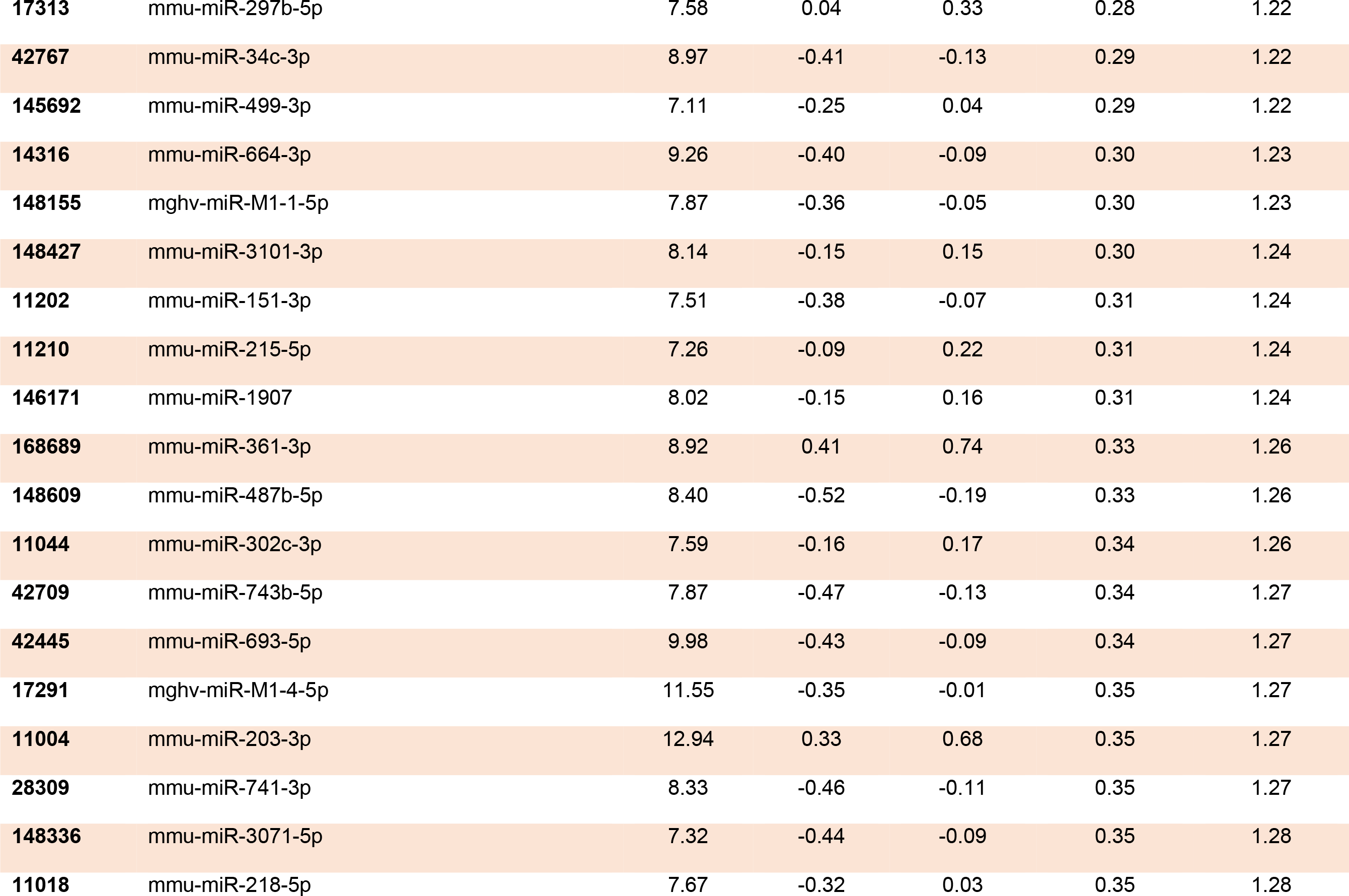
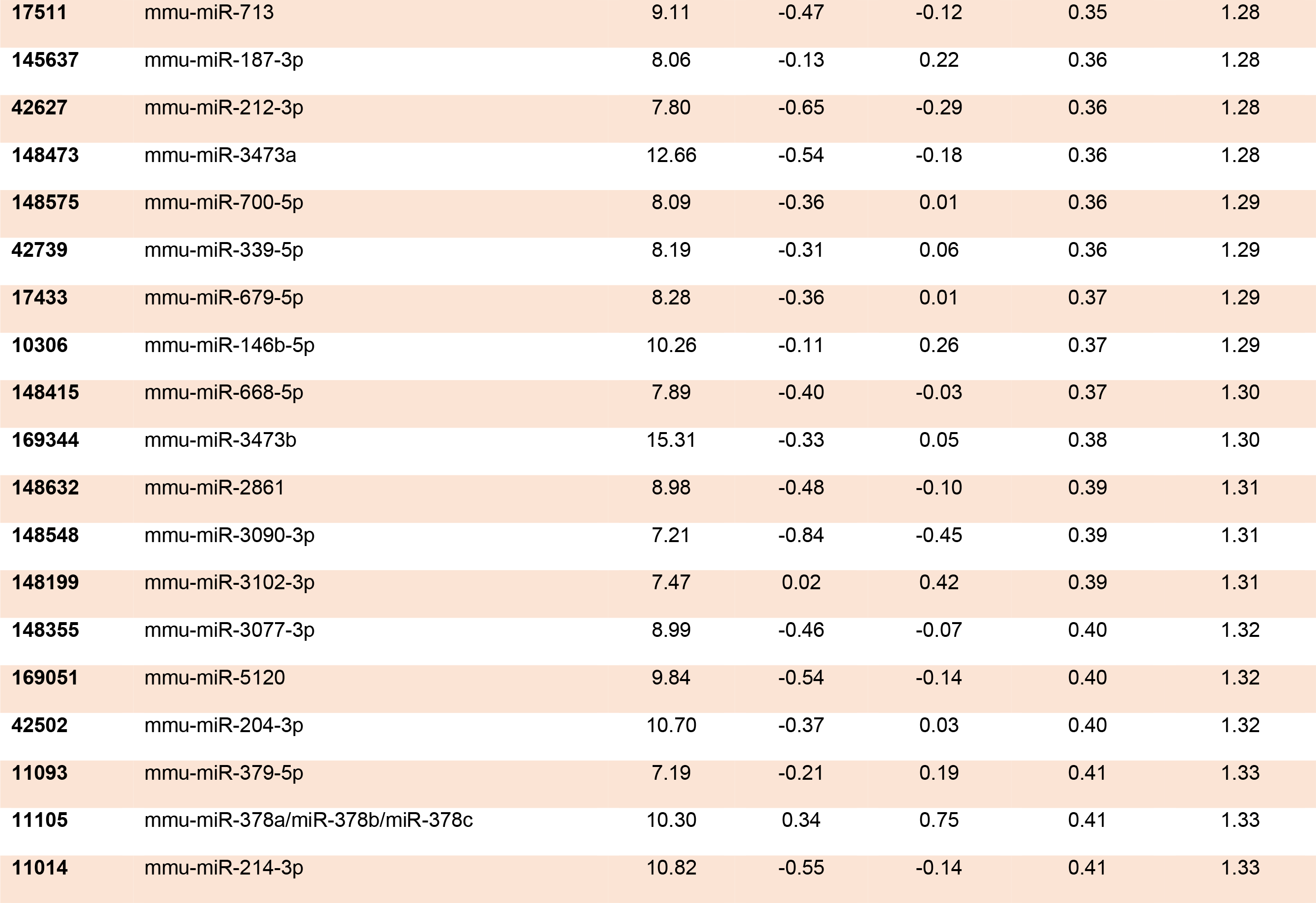
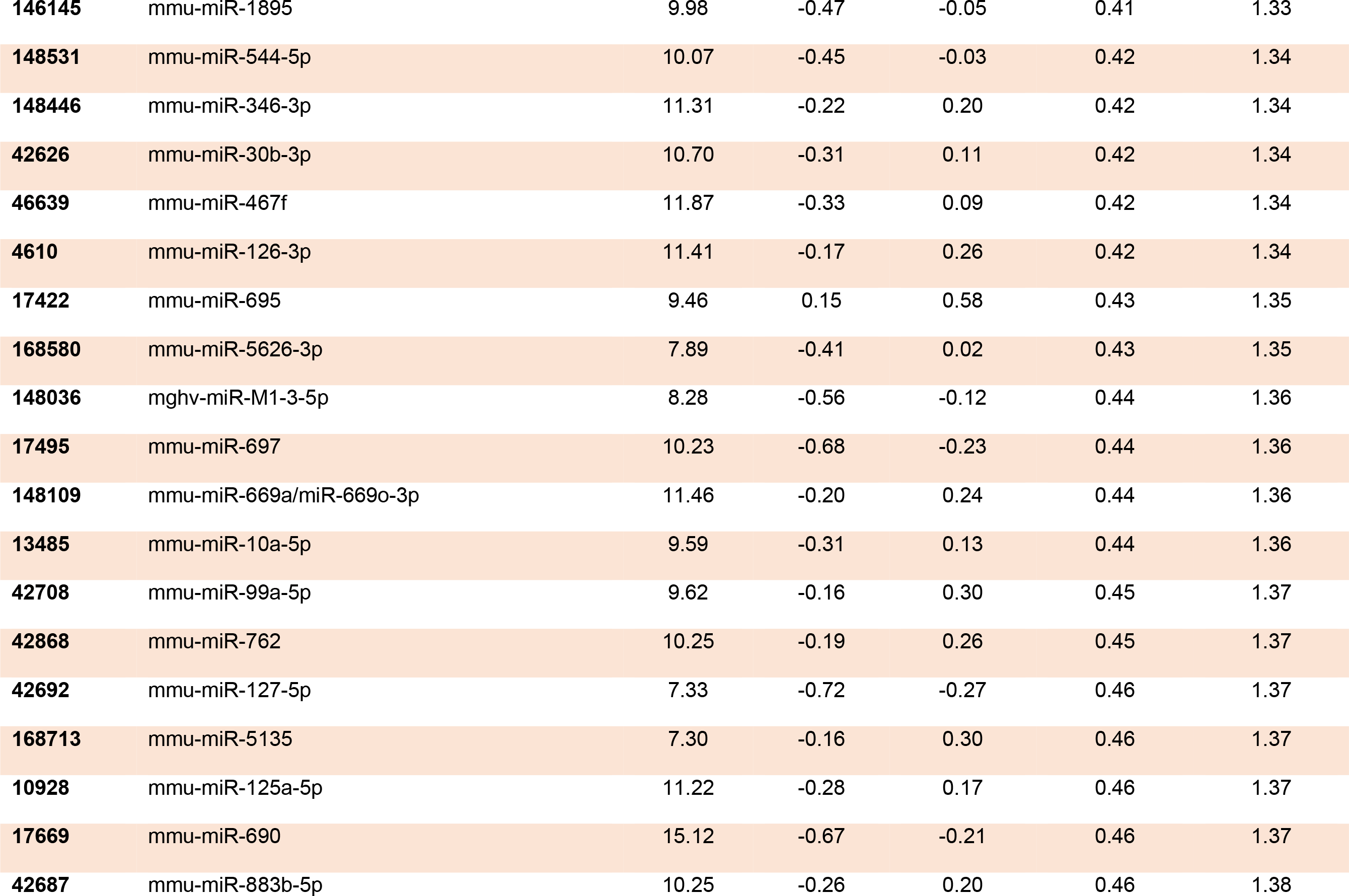
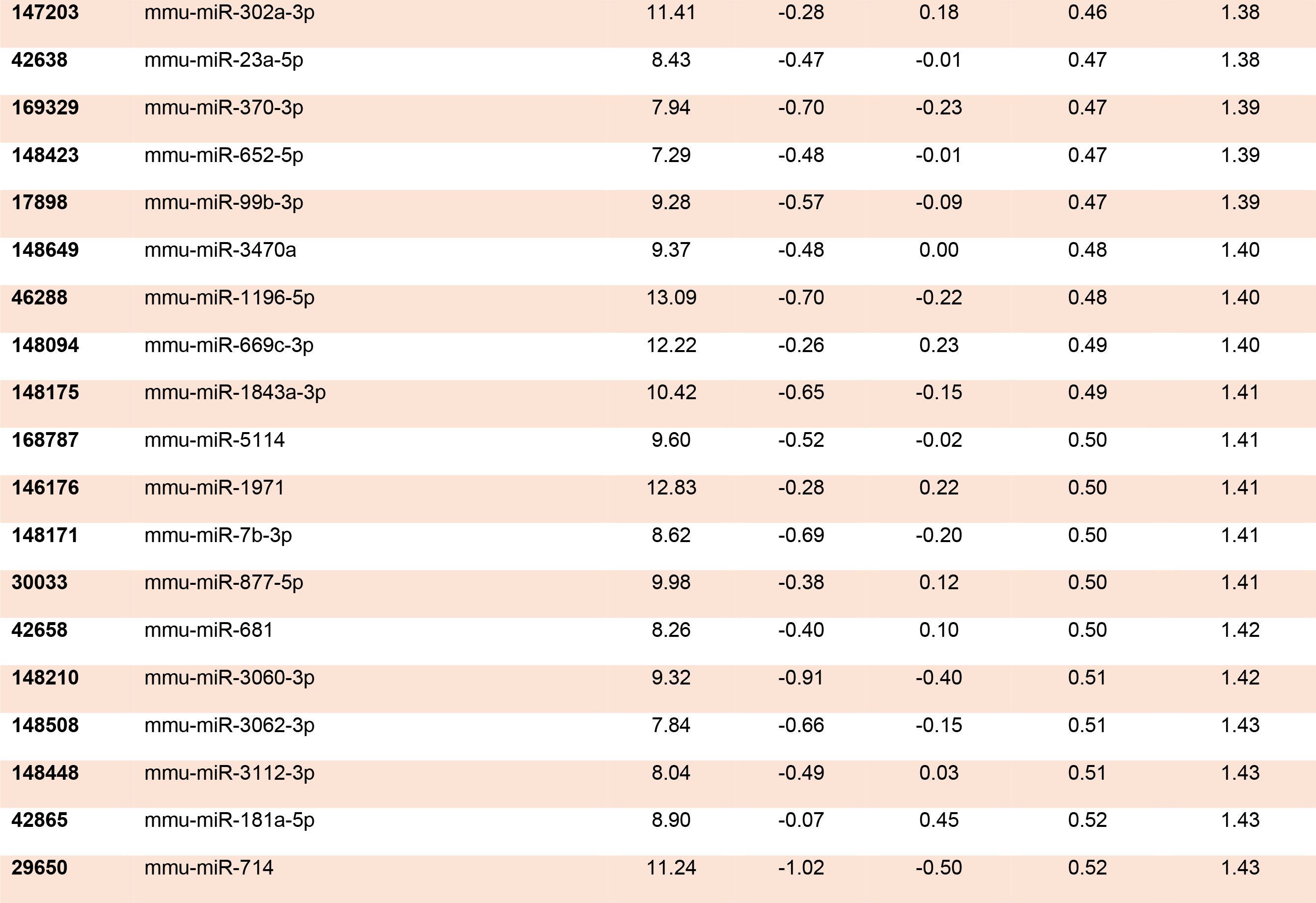
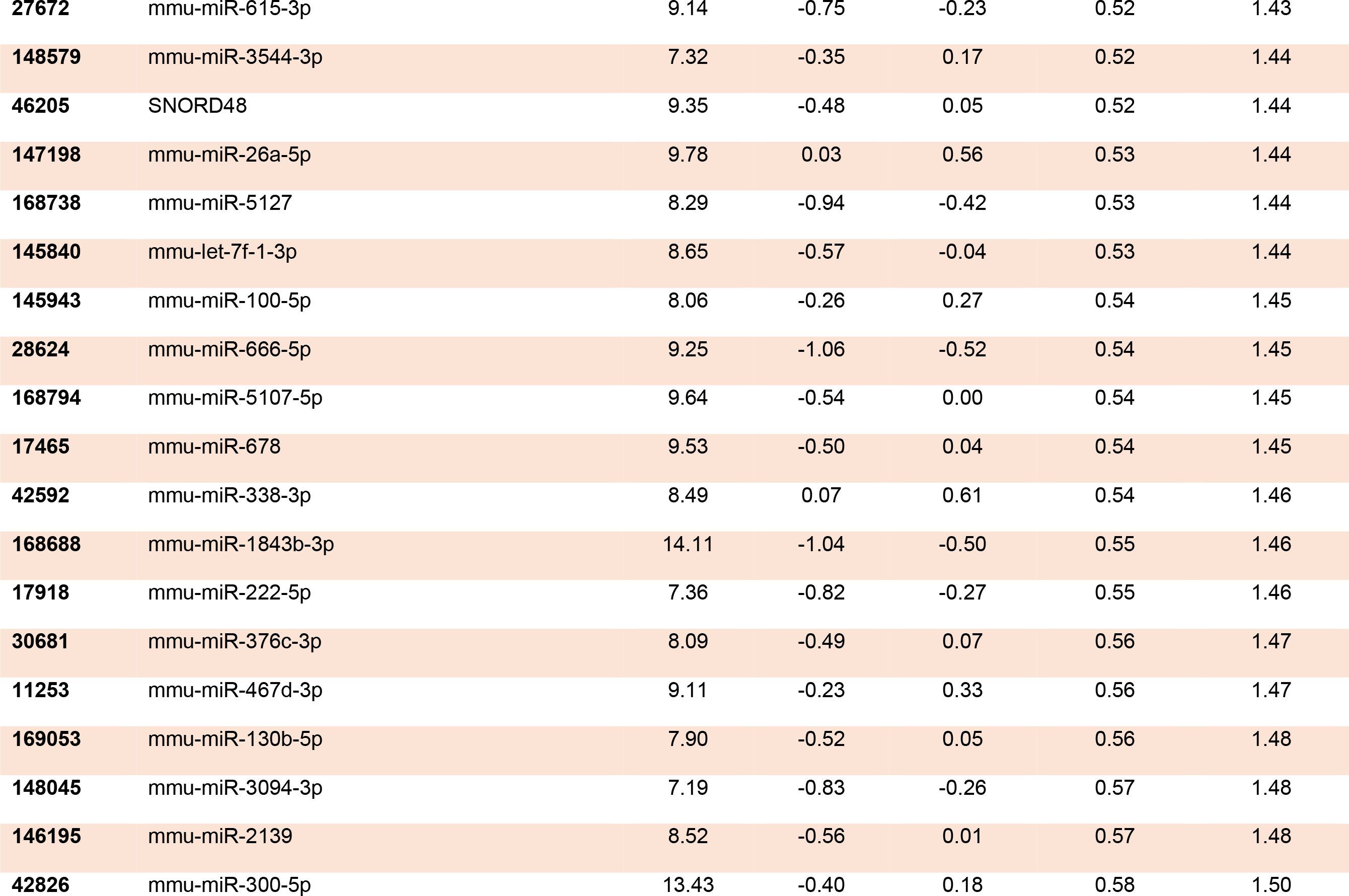
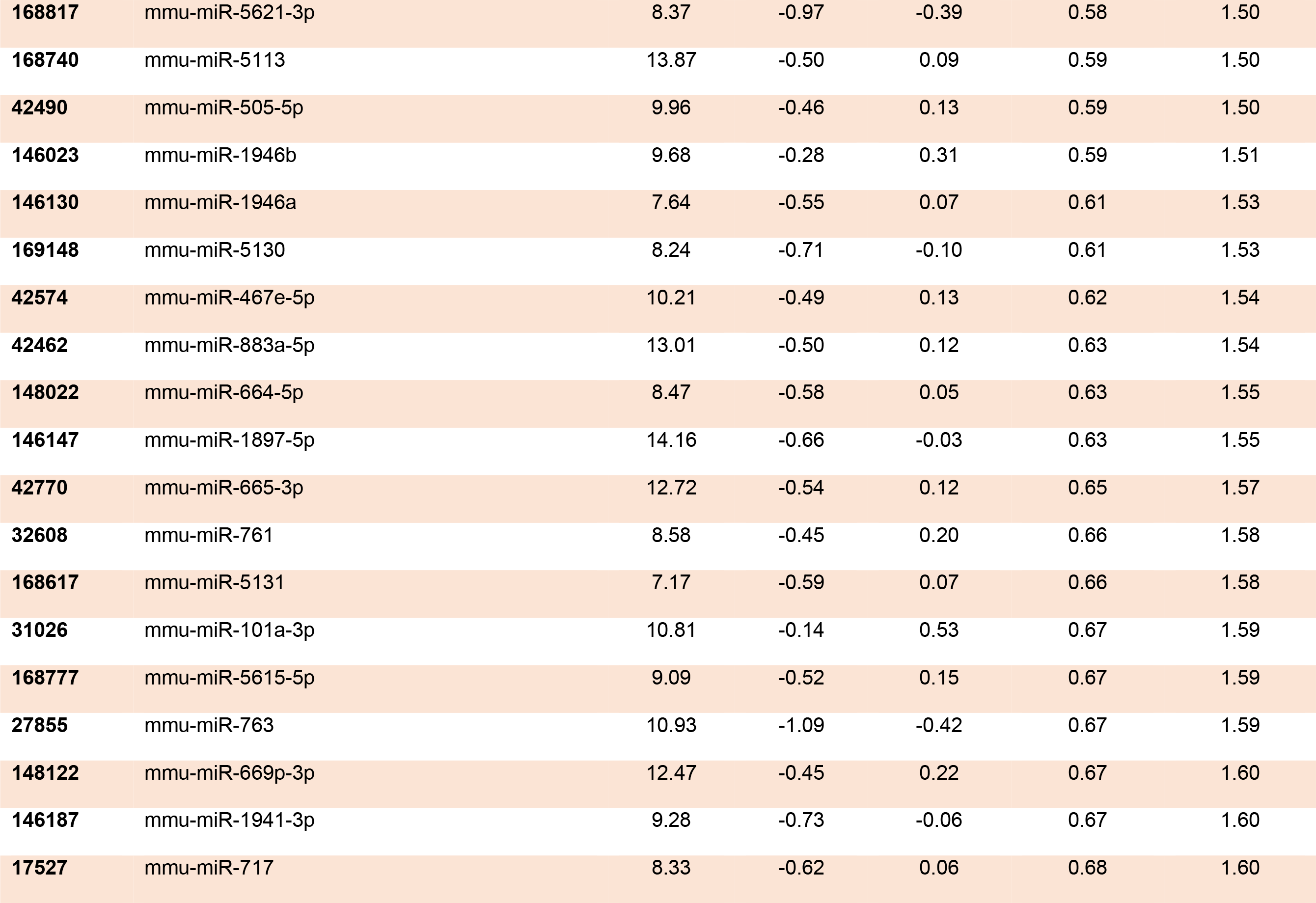
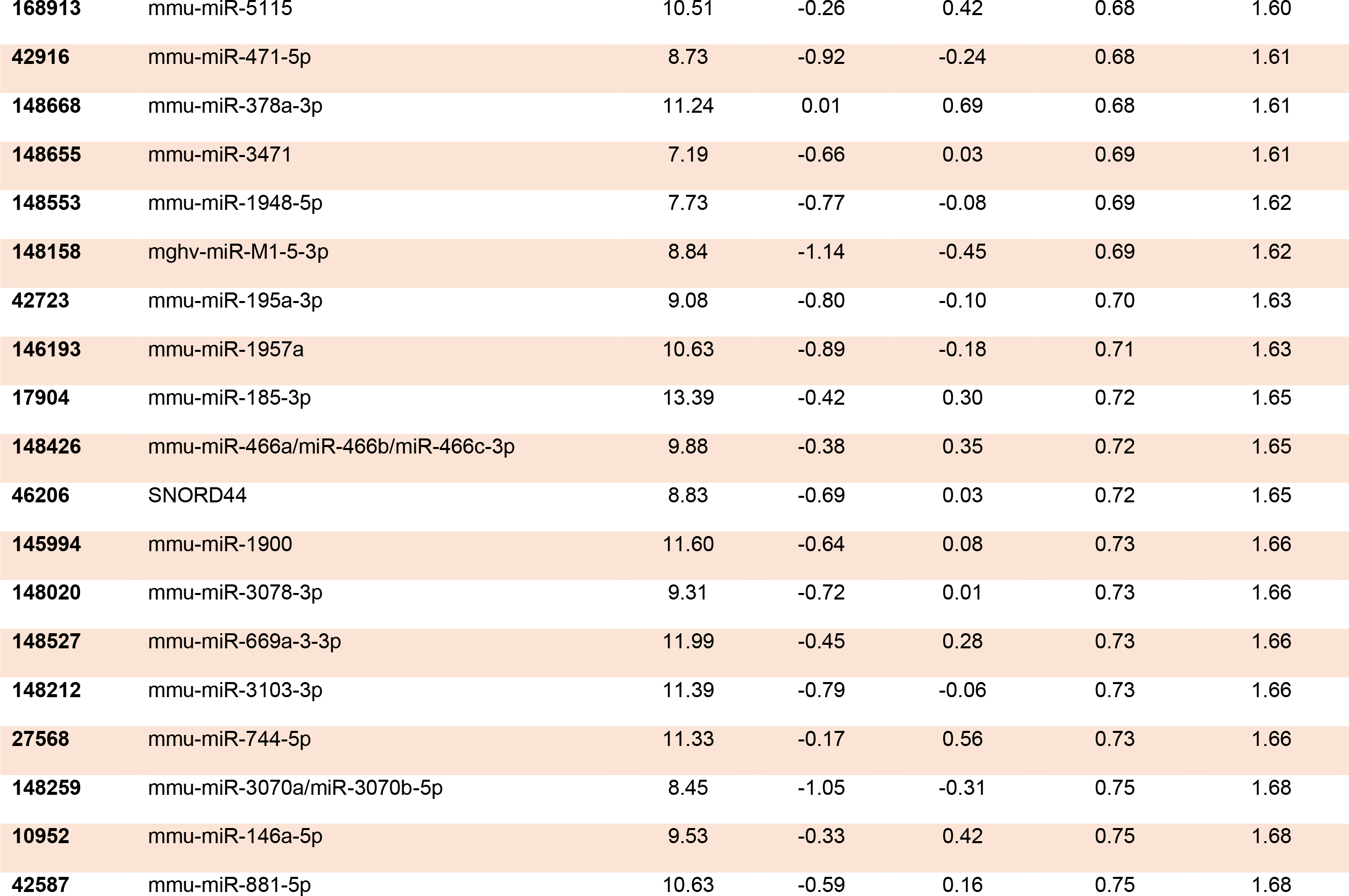
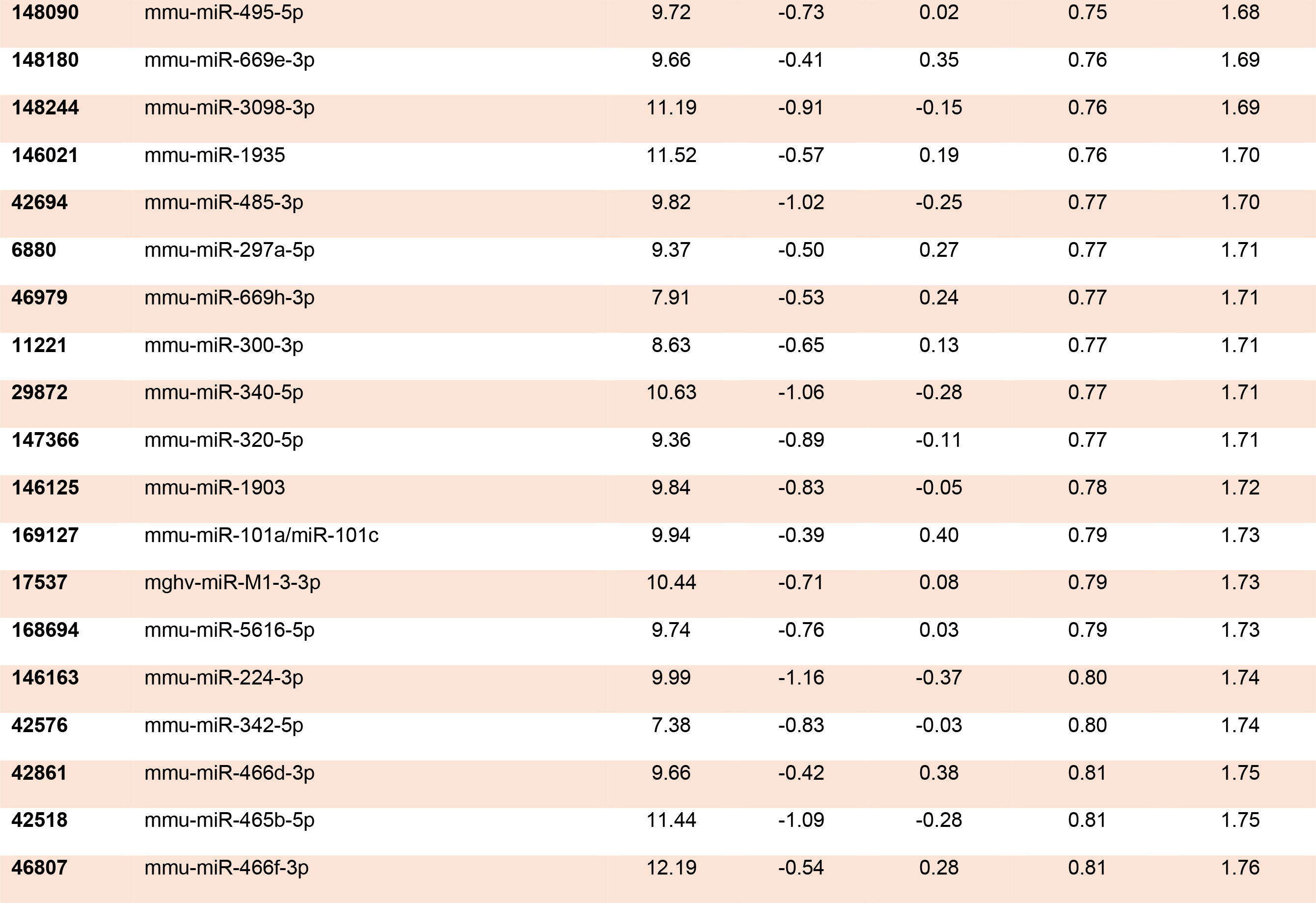
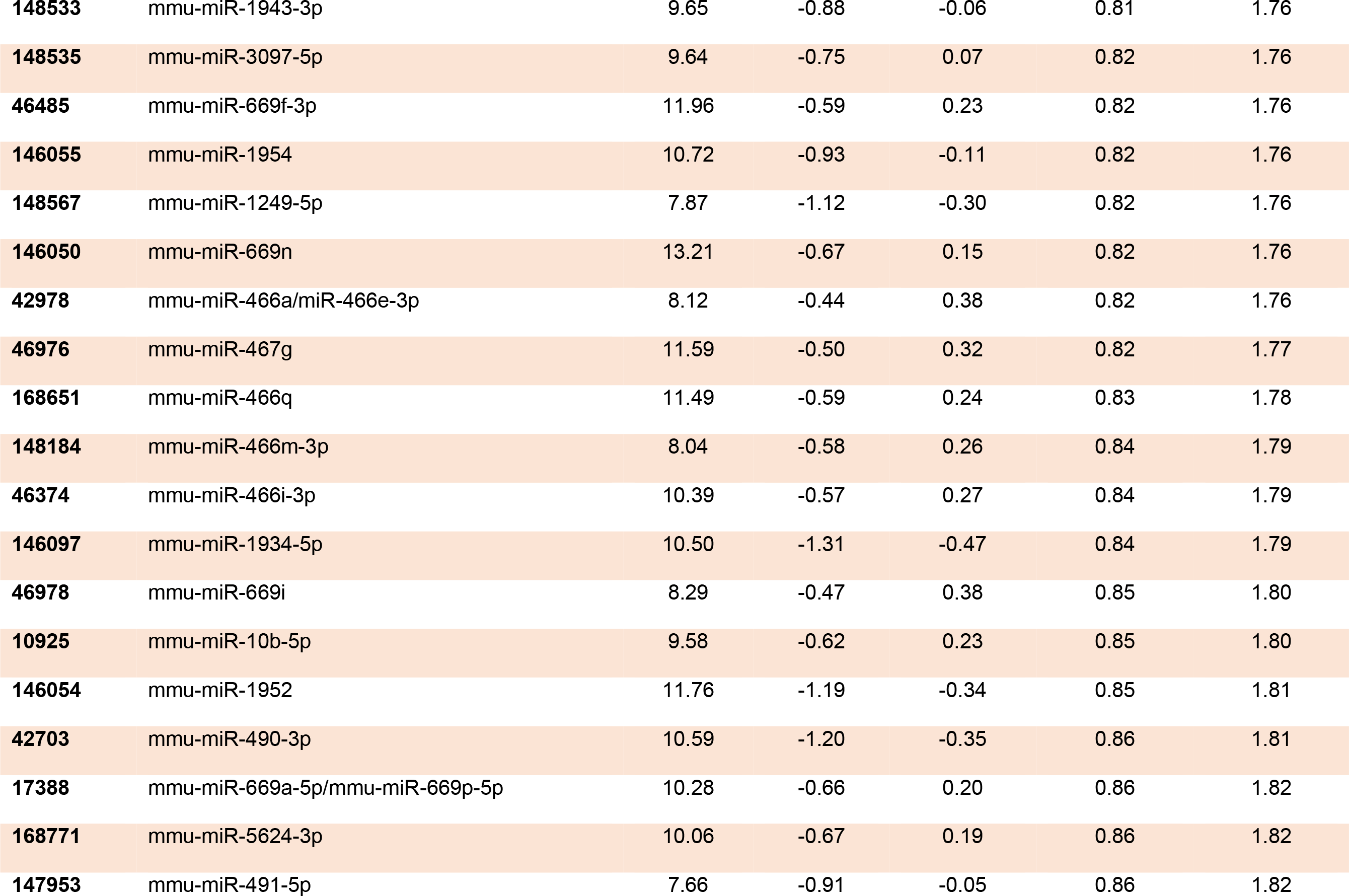
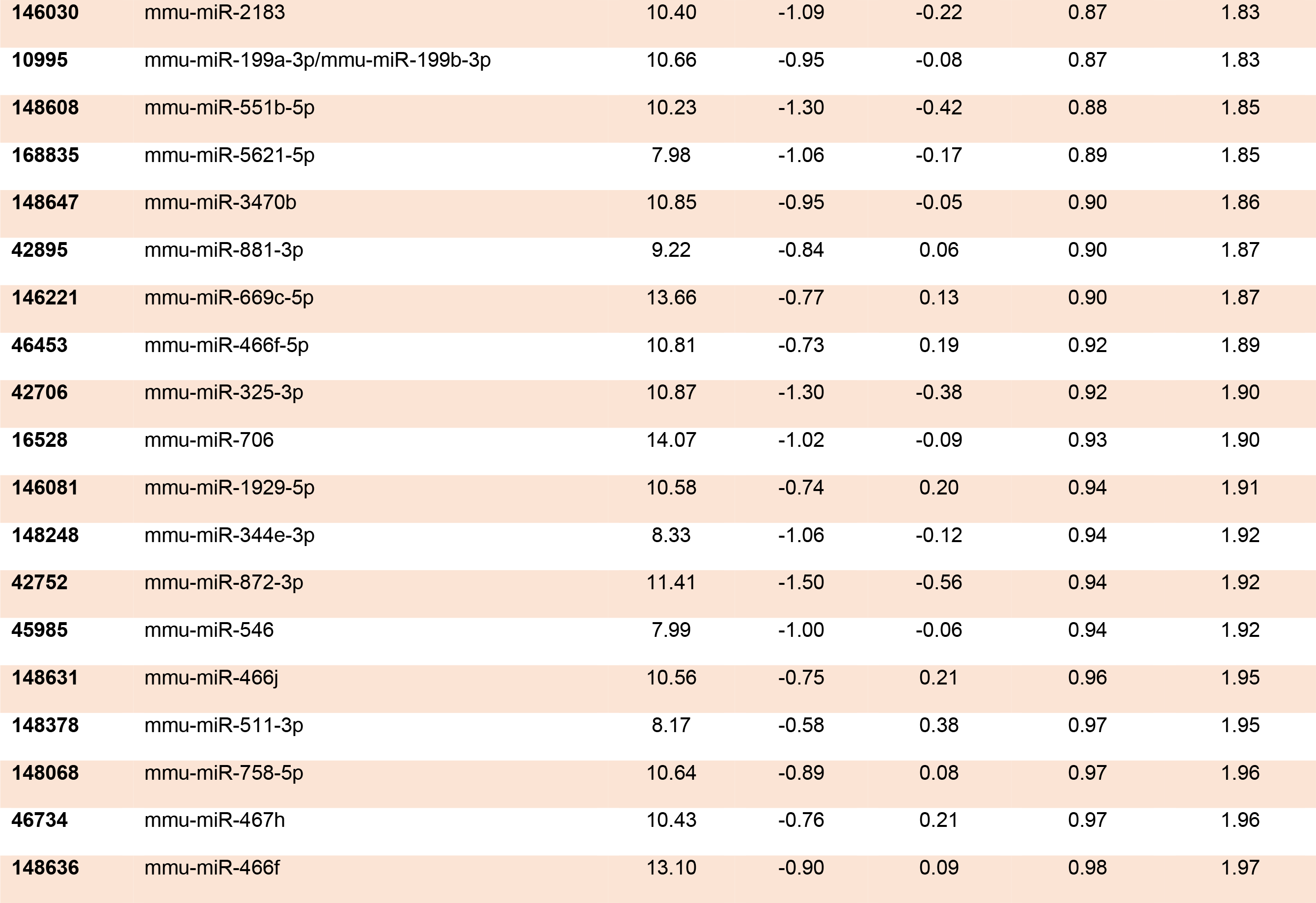
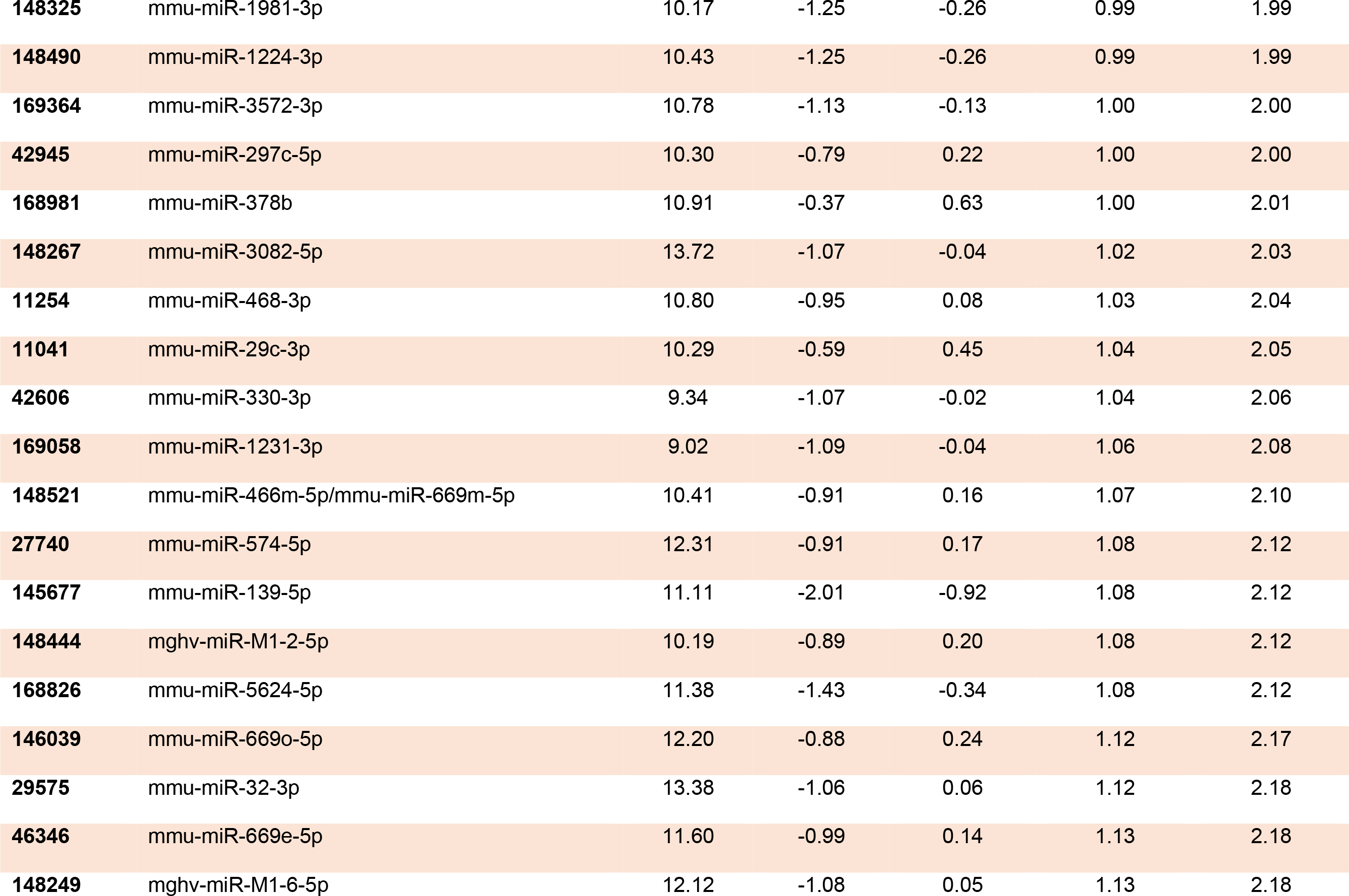
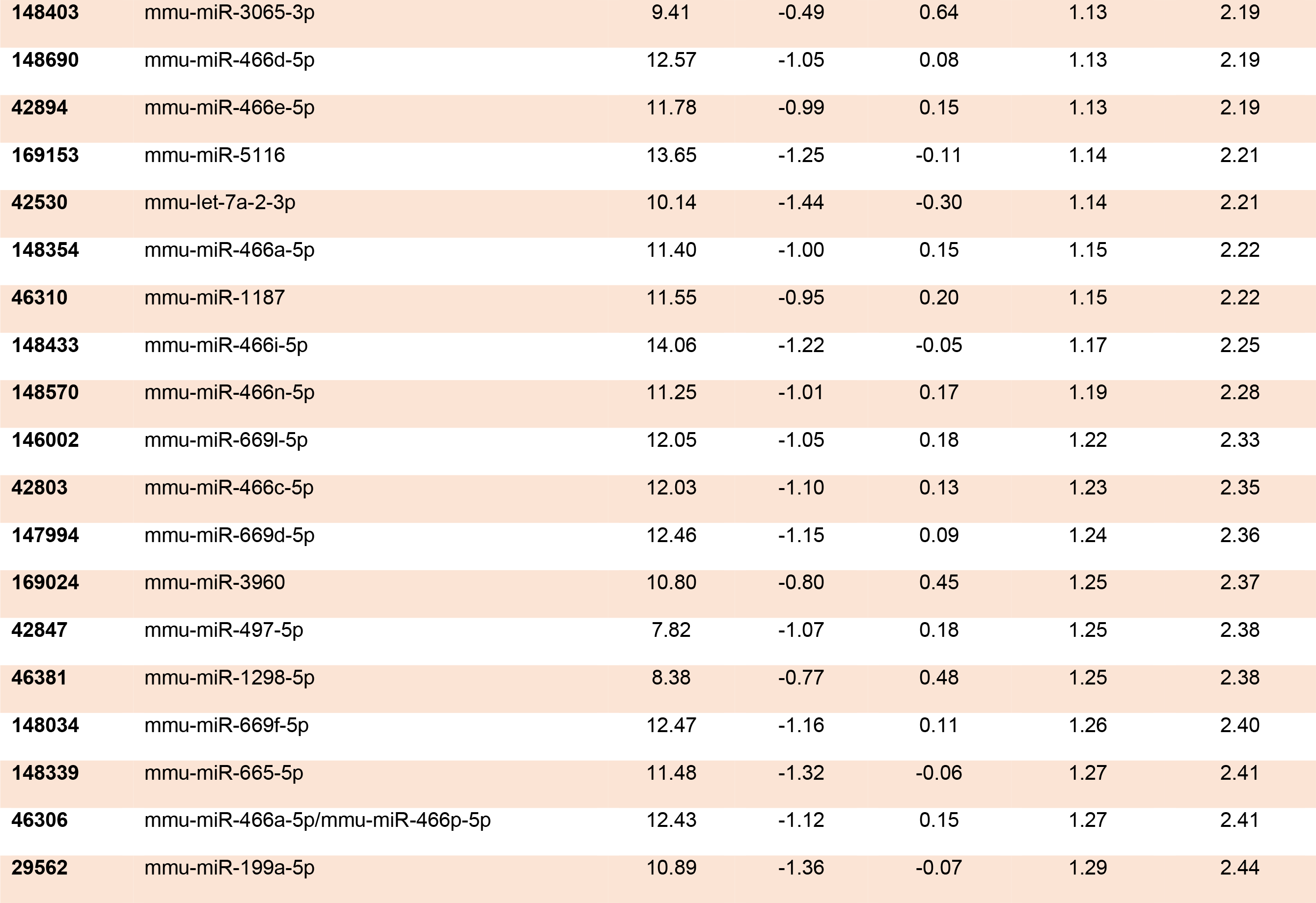
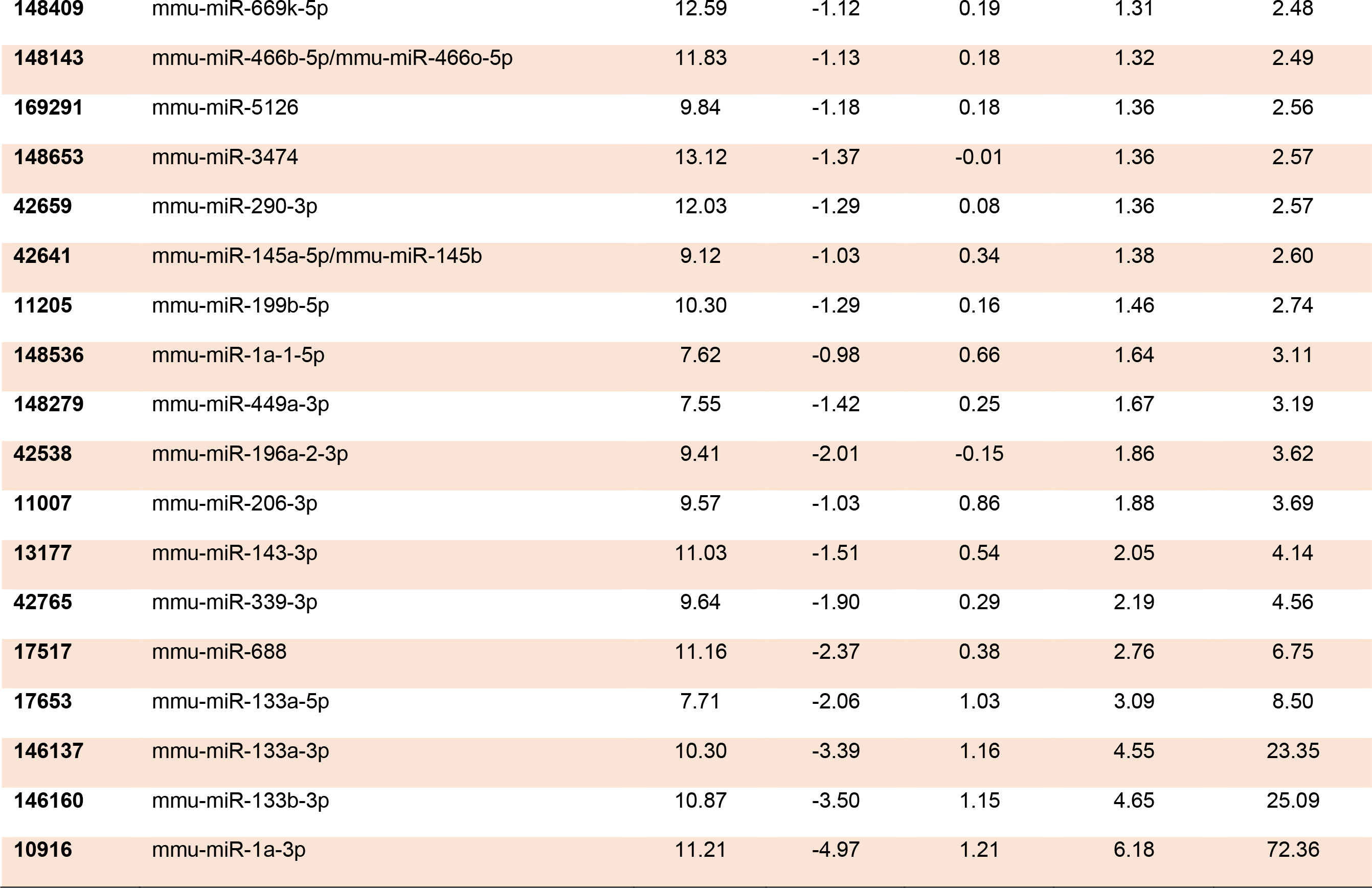
Fold change expression of up-regulated miRNAs in UVB-induced tumors (UT). Data were compared with non-UVB exposed normal skin (CS).

### 2.4. Effect of mimic of miR-15b and an inhibitor of miR-133a on UVB-induced immune suppression of the CHS response

UVB-induced immunosuppression has been considered a key mechanism in photocarcinogenesis [21, 22]. Therefore, to identify the role of miRNAs altered in UVB-induced tumors, the two miRNAs (miR-15b and miR-133a) were randomly selected for their role in immunosuppression in the CHS study. As shown in **Fig. 4A and 4B**, UVB exposure suppresses the CHS response of DNFB upto 86.45% (p<0.0018; bar 3) compared with the non-UVB exposed group (bar 2). Intraperitoneal injections of mimic of downregulated miR-15b (30nM) block UVB-induced immunosuppression upto 42.05% (p<0.0218; bar 4, **Fig. 4A**), while treatment with an inhibitor of upregulated miR-133a (20 nM) blocks UVB mediated immune suppression upto 69.89% (p<0.0041; bar 4, **Fig. 4B**). These observations revealed that maintenance of UVB mediated miRNAs protect mice from UVB induced immune suppression.

**Figure 4.**
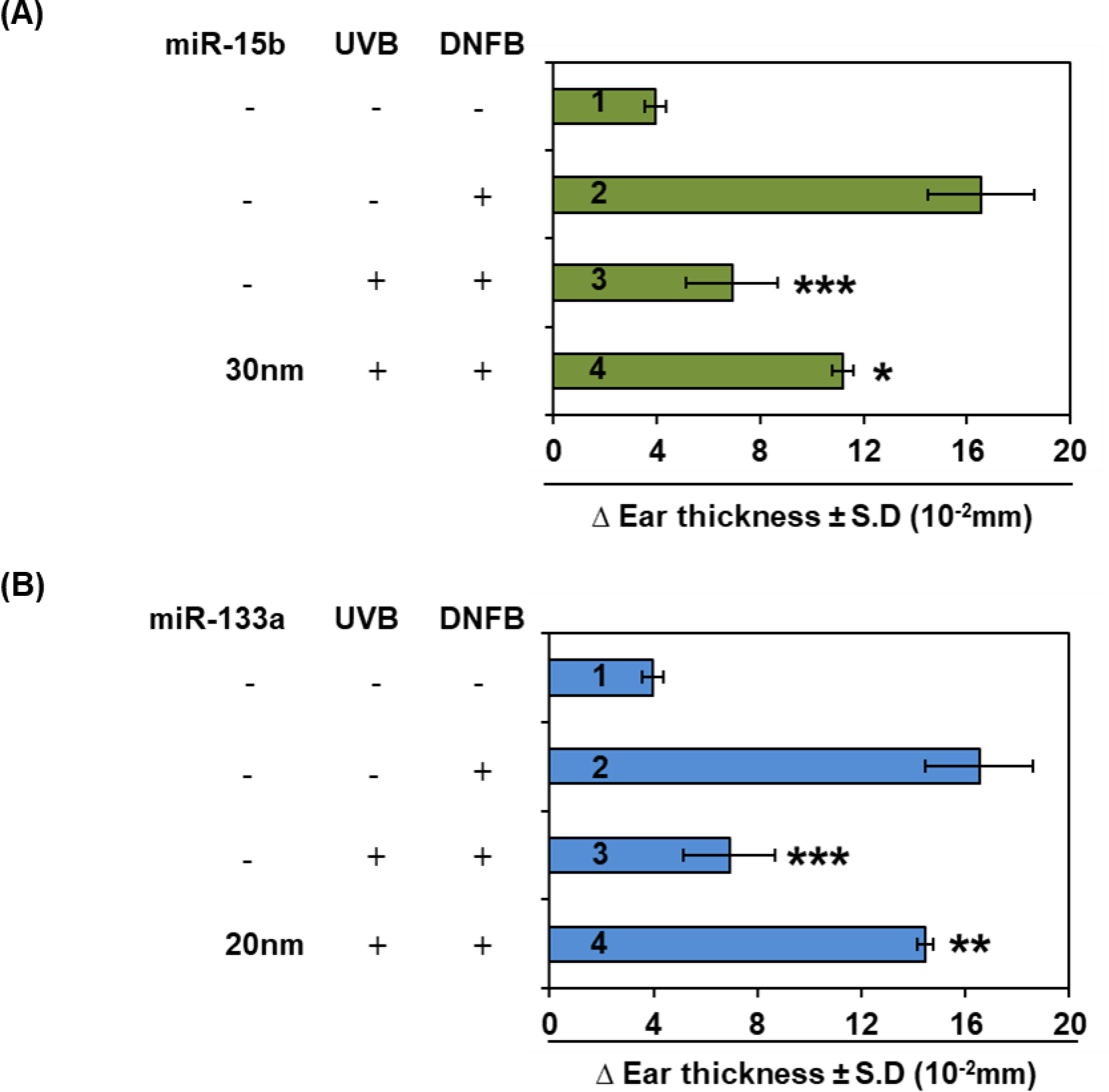
The effect of intraperitoneal injections of mimic of miR-15b **(A)** and inhibitor of miR- 133a **(B)** on UVB induced immunosuppression. The mice were treated with mimic/inhibitor 30 mins prior to UVB exposure. The data is presented as change in mean ear thickness ± S.E. n = 4, Statistical differences; *p<0.05, **p<0.01, ***p<0.001.

## Discussion

Exposure to UV radiation is the principal cause of non-melanoma skin cancer, the most prevalent human cancer, and is also associated with the induction of malignant melanoma. The global incidence of both types of skin cancer (non-melanoma and melanoma) has been increasing over the past few years. Approximately 2-3 million non-melanoma and 132,000 melanoma skin cancers occur worldwide yearly. According to the skin care foundation, one in every three cancers is skin cancer diagnosed globally [23]. UV radiation is a complete carcinogen, which can initiate skin cancer followed by promotion and progression. Exposure to UV radiation of the skin causes inflammation, apoptosis, DNA damage, oxidative stress, immunosuppression, and premature aging of the skin [24, 25].

Cancer formation is the combined interaction of tumor suppressors and oncogenes. Although several genes have been identified in human and animal models, the mechanism of cancer formation is yet to be determined. A recent study demonstrated that more than 50% of miRNA genes are located in cancer-associated genomic regions or fragile sites [26], suggesting that miRNAs may play a more critical role in the pathogenesis of a limited range of human cancers. Various studies have shown that miRNAs play essential roles in DNA methylation, cell proliferation, differentiation, angiogenesis, cell survival, and activation of several molecular pathways in cancers, such as miRNA-106b regulating cell survival pathways and enhancing cell proliferation and tumor of melanoma cells by targeting p21 expression [27]. Several other miRNAs, such as miRNA-17, miRNA-18, miRNA-19, miRNA-21, miRNA-29c, etc., have been reported for their roles in non-melanoma skin cancer [28, 29].

In the miRNA profiling of UVB tumor tissues, we observed that the expression levels of several miRNAs were altered either upregulated or downregulated, and they may have an important role in photocarcinogenesis. When cells exhibit abnormal growth and loss of apoptosis function, it usually results in cancer formation. Recent studies indicate that miRNA regulates cell growth and apoptosis [30, 31]. For example, *miR-15* and *miR-16* induce apoptosis by targeting anti-apoptotic gene B cell lymphoma 2 (*BCl2*) mRNA [32], which is a key player in many types of human cancers, including skin cancer. Our profiling demonstrated that expression levels of *miR-15* and *miR-*16 were upregulated, suggesting that these miRNAs may have been involved in UVB-induced tumor formation through the regulation of cell growth and apoptosis. The expression of miRNA cluster miRNA-17-92 was also observed to be upregulated in UVB tumors. Recent studies reported that the expression of miRNA cluster miR-17–92 is remarkably increased in several other cancers, including lung cancer [33], by targeting two tumor suppressor genes, *PTEN* and *RB1* [33]. *PTEN* promotes apoptosis through the P13K-Akt- PKB pathway [34]. In our results, we also observed that these miRNAs were upregulated in UVB-induced tumors and provided evidence for their role in photocarcinogenesis. Cancer is the manifestation of genetic and epigenetic events. Exposure to UVB radiation causes DNA hypermethylation and silenced tumor suppressor genes to promote tumor growth. Recent studies indicate that loss of miRNA-29s expression has a role in DNA methylation by targeting DNA methyltransferases and TET enzymes [35, 36]. Our profiling results demonstrated that the levels of miRNA-29s were down regulated in UVB tumors.

Briefly, miRNAs expression profiling helps to identify a new range of miRNAs that regulate several biological processes and may involve in various diseases, including carcinogenesis and provides a better platform for distinguishing cancer tissues from normal tissues. Our observations suggest that miRNAs can be used as biomarkers and a powerful diagnostic tool for detecting cancers.

In conclusion, for the first time, we have shown that such an expression profiling approach is a suitable and effective solution for identifying aberrant miRNAs involved in cancer progression caused by UV radiation. The results of this miRNAs expression profiling in UVB-induced tumors will provide new insights into discovering potential biomarkers in photocarcinogenesis. Further mechanistic and external validation studies are needed for their clinical significance and role in the development of skin cancer.

## 4. Material and methods

### 4.1. Animals

SKH-1 hairless mice (6-7 weeks old) were purchased from Charles River Laboratory (Wilmington, MA). The mice were kept for at least one week in our animal resource facility before use in experiments. Mice were maintained under standard conditions of a 12-h dark/12-h light cycle, a temperature of 24 ± 2°C, and relative humidity of 50 ± 10%. The animal study was approved by the Institutional Animal Care and Use Committee (IACUC) of the University of Alabama at Birmingham.

### 4.2. UVB irradiation and photocarcinogenesis protocol

The mice were exposed to UVB radiation, as described previously [37]. Briefly, the dorsal skin of SKH-1 hairless mice was exposed to UVB radiation from a band of four FS24T1 UVB lamps (Daavlin, UVA/UVB Research Irradiation Unit, Bryan, OH) equipped with a regulator for UVB dosa. Under the standard photocarcinogenesis protocol, mice were UVB irradiated (180 mJ/cm^2^; 3X/week) for upto 24 weeks. At the end of the study, mice from both cohorts were humanly euthanized, and samples were collected and stored at -80^0^C.

### 4.3. Tissue collection and RNA isolation

At the end of the photocarcinogenesis experiment, tumor and skin tissues were harvested after the euthanization of experimental mice. Total RNAs, including small ones, were extracted from the skin/tumor tissues using the Qiagen miRNeasy® mini kit. Briefly, a small portion of tissue (10mg) was lysed in 700µL Qiazol lysis reagent using a tissue lyzer with one 5mm stainless steel bead. The tissue lysate was transferred to a new tube with 140 µL chloroform, mixed, incubated for 2 min at room temperature, and centrifuged at 12,000 x g for 15 min at 4^0^C. The upper aqueous phase was separated, and 525µL of 100% ethanol was added. The contents were mixed gently, transferred into RNeasy mini spin column in a collection tube, and centrifuged at 8,000 x g for 15 sec at room temperature. The RNeasy mini spin column was rinsed with 700 µL RWT buffer and centrifuged at 8,000 x g for 15 sec at room temperature, followed by another rinse with 500 µL RPE buffer and centrifuged at 8,000 x g for 15 sec at room temperature. The rinse step with RPE buffer was repeated two times. After centrifugation, the flow-through was discarded, RNeasy mini spin column was transferred to a new collection tube, and the lid was left uncapped for 1 min to allow the column to dry. Total RNA was eluted by adding 50 µL of RNase-free water to the membrane of the RNeasy mini spin column and incubating for 1 min before centrifugation at room temperature. The RNAs, including small RNAs, were stored at -80^0^C until used.

### 4.4. miRNAs array profiling

All microRNAs array was conducted at Exiqon Services, Denmark. An Agilent 2100 Bioanalyzer profile verified the quality of the total RNA. 750 ng total RNA from both sample and reference was labeled with Hy3™ and Hy5™ fluorescent labels, respectively, using the miRCURY LNA™ microRNA Hi-Power labeling kit, Hy3™/Hy5™ (Exiqon, Denmark) following the procedure described by the manufacturer. The Hy3™ labeled samples and a Hy5™ labeled reference RNA sample was mixed pairwise and hybridized to the miRCURY LNA™ microRNA array 7th Gen (Exiqon, Denmark), which contains capture probes targeting all microRNAs for human, mouse, and rat registered in the miRBASE 18.0. The hybridization was performed according to the miRCURY LNA™ microRNA array instruction manual using a Tecan HS4800™ hybridization station (Tecan, Austria). After hybridization, the microarray slides were scanned and stored in an ozone-free environment (ozone level below 2.0 ppb) to prevent potential bleaching of the fluorescent dyes. The miRCURY LNA™ microRNA array slides were scanned using the Agilent G2565BA Microarray Scanner System (Agilent Technologies, Inc., USA), and the image analysis was carried out using the ImaGene® 9 (miRCURY LNA™ microRNA Array Analysis Software, Exiqon, Denmark). The quantified signals were normalized using the global LOWESS (Locally Weighted Scatterplot Smoothing) regression algorithm and background corrected [38]. As per the recommendations provided by Exiqon, smaller fold changes (≤1.0 fold) may tend to be relatively more affected by technical variance, and such changes could be associated with the increased risk of false-positive signals. Therefore the cut-off value was ≤1.0 fold change in our study. The data for miRNAs expression showing ≤1.0 fold change were eliminated during comparison within the groups.

### 4.5. Contact hypersensitivity (CHS) assay

The effect of miRNA-15b and miRNA-133a mimic/inhibitor on UVB-induced immune suppression in mice was assessed using the contact hypersensitivity model described previously [21, 22]. Briefly, dorsal skin-shaved mice were exposed to UVB radiation (150 mJ/cm^2^) on four consecutive days. During the UVB exposure, ears were protected from UV irradiation by covering. After 24 hours of last UVB exposure, mice were sensitized with skin contact sensitizer 2, 4-dinitrofluorobenzene (DNFB) by topical application [(0.5% in 25 µl of acetone: olive oil mixture (4:1, v/v)]. After five days, CHS response was elicited by treating with 20 µl of 0.2% DNFB (ears). The thickness of ear skin was measured 24 h after the challenge using an engineer’s micrometer (Mitutoyo, Tokyo, Japan). The CHS response was calculated by comparing the ear thickness before the challenge. To determine the effect of miRNAs on UVB-induced immune suppression, mice were administered with the mimic of miRNA-15b (30nM; i.p) and an inhibitor of miRNA-133a (20nM; i.p.). The experimental mice were treated with miRNAs mimic/inhibitor daily, 30 mins before UVB exposure.

### 4.6. Statistical Analysis

Data were evaluated for outliers and adherence to a normal distribution using GraphPad Prism software (San Diego, CA, USA), version 8.1. Statistical significance of normally and non-normally distributed data were assessed via one-way ANOVA and Tukey’s multiple comparison test, respectively, with α = 0.05.

## Author Contributions

A.A., V.K., H.F., V.K.S, and R.P. conceived the study and participated in the design. R.P. participated in sample collection and CHS study. All authors wrote, edited, and consented to the published version of the manuscript.

## Data Availability Statement

The original data presented in the study are available on request from the corresponding author.

